# Rapid *in*-*situ* diversification rates in Rhamnaceae explain the parallel evolution of high diversity in temperate biomes from global to local scales

**DOI:** 10.1101/2023.08.26.554607

**Authors:** Qin Tian, Gregory W. Stull, Jürgen Kellermann, Diego Medan, Francis J. Nge, Shuiyin Liu, Heather R. Kates, Douglas E. Soltis, Pamela S. Soltis, Robert P. Guralnick, Ryan A. Folk, Renske E. Onstein, Ting-Shuang Yi

## Abstract

- The macroevolutionary processes that have shaped biodiversity across the temperate realm remain poorly understood and may have resulted from evolutionary dynamics related to diversification rates, dispersal rates, and colonization times, closely coupled with Cenozoic climate change.
- We integrated phylogenomic, environmental ordination, and macroevolutionary analyses for the cosmopolitan angiosperm family Rhamnaceae to disentangle the evolutionary processes that have contributed to high species diversity within and across temperate biomes.
- Our results show independent colonization of environmentally similar but geographically separated temperate regions mainly during the Oligocene, consistent with the global expansion of temperate biomes. High global, regional, and local temperate diversity was the result of high *in*-*situ* diversification rates, rather than high immigration rates or accumulation time, except for Southern China, which was colonized much earlier than other regions. The relatively common lineage dispersals out of temperate hotspots highlights strong source-sink dynamics across the cosmopolitan distribution of Rhamnaceae.
- The proliferation of temperate environments since the Oligocene may have provided the ecological opportunity for rapid *in*-*situ* diversification of Rhamnaceae across the temperate realm. Our study illustrates the importance of high *in*-*situ* diversification rates for the establishment of modern temperate biomes and biodiversity hotspots across spatial scales.

## Introduction

Understanding the historical processes responsible for heterogeneity in the distribution of species richness across the globe is a major goal in evolutionary biology (Schluter & Pennell, 2017). The most prominent diversity pattern at the global scale is the tendency for species richness to increase towards the equator, i.e., the latitudinal diversity gradient (Hillebrand, 2004; Jablonski *et al*., 2006; Mittelbach *et al*., 2007). However, deviations from this global pattern, such as the high species richness and endemism in temperate biomes (i.e., bimodal latitudinal diversity gradient) rather than tropical biomes, remain puzzling (Orr *et al*., 2021). Indeed, one third of the world’s biodiversity hotspots (Myers *et al*., 2000) – regions that contain high levels of plant species richness and endemism yet are under threat from human activities – are located in temperate zones (Igea & Tanentzap, 2019), such as the five Mediterranean-type ecosystems (MTEs, i.e., California Floristic Region, Mediterranean Basin, Cape Floristic Province, Southwest Australia, Chilean Winter Rainfall-Valdivian Forests; Rundel *et al*., 2016) and the Northern-Hemisphere mountain hotspots (e.g., Hengduan Mountains, Himalayas; Xing & Ree, 2017). A series of studies has clarified patterns of high species richness and endemism in specific temperate areas, such as mountains, through uplift-driven diversification (Hughes, 2017; Zhang *et al*., 2021). However, whether high diversification rates characterize temperate biodiversity hotspots more generally remains unclear due to the lack of a global analysis focused on temperate diversification. A global perspective is needed to understand whether temperate biodiversity is the result of similar drivers, such as climatic changes at global scales (e.g., global cooling and aridification since the Eocene-Oligocene boundary; Zachos *et al*., 2001) and evolutionary processes (e.g., rapid speciation), or whether each region has a unique set of diversity drivers.

Current species diversity in temperate regions may be the outcome of three main evolutionary processes. First, *in*-*situ* diversification rates – the balance between speciation and extinction within temperate regions – may have contributed to extant diversity. Indeed, many plant evolutionary radiations characterized by high diversification rates are thought to have occurred entirely in temperate biomes, such as the *Phylica* (Rhamnaceae) and *Moraea* (Iridaceae) radiations in the Cape flora of southern Africa (Goldblatt *et al*., 2002), and the *Dianthus* (Caryophyllaceae), *Tragopogon* (Asteraceae), and *Cistus* (Cistaceae) radiations in the Mediterranean Basin (Guzman *et al*., 2009; Valente *et al*., 2010; Bell *et al*., 2012; Rundel *et al*., 2016). Second, differences in historic rates of lineage dispersal can result in some temperate areas acting as ‘sources’, i.e., providing frequent dispersal of lineages emigrating to other regions where climatic conditions are suitable, and others as ‘sinks’, with high dispersal rates of lineages immigrating into the region, thereby increasing overall species richness. Immigration has been shown to play a significant role in the formation of regional biodiversity in the Hengduan Mountains and Himalayas (Xing & Ree, 2017; Ding *et al*., 2020). Lastly, the time-for-speciation hypothesis (Stephens & Wiens, 2003) states that gradual diversification over time may lead to the build-up of species richness in a region, predicting that dispersal and diversification regimes do not differ among regions but instead regions that were colonized early harbor higher species richness than regions colonized later, either as a result of differences in biome or region age or differences in colonization opportunities. Time-for-speciation is often invoked to explain high diversity in the tropics, because tropical biomes are generally thought to be older than temperate ones, and may thus have accumulated diversity over longer periods of time (Mittelbach *et al*., 2007; Pontarp *et al*., 2019). Overall, contrasting mechanisms have been invoked to explain the distribution of biodiversity across individual temperate areas, but we lack global clarity on which processes are most important.

Macroevolutionary processes of lineage diversification, dispersal, and diversity accumulation reflect a background of dynamic environmental and geological change. For example, although hot and humid rainforest biomes probably date back to the Cretaceous (ca. 100 Ma) at middle paleolatitudes (Morley, 2000), climate cooling and aridification since the Eocene-Oligocene boundary (ca. 34 Ma), has led to the global proliferation of temperate biomes (Zachos *et al*., 2001; Palazzesi *et al*., 2022). Expansion of temperate biomes is thought to have provided ecological opportunities for lineages to colonize and diversify (Simpson, 1953; Donoghue, 2008; Stroud & Losos, 2016). Indeed, many temperate-adapted lineages evolved and diversified around and after Oligocene, such as drought- and cold-adapted C_4_ grasses (Poaceae), succulents (e.g., Aizoaceae, Cactaceae), and orchids (Orchidoideae), leading to the subsequent spread of grasslands and deserts (Spriggs *et al*., 2014; Palazzesi *et al*., 2022; Thompson *et al*., 2023a; Thompson *et al*., 2023b). In most of these cases, it is thought that lineages already possessed traits needed for colonizing a new region, and that the niche was largely conserved as lineages radiated after arrival (Wiens & Donoghue, 2004; Donoghue, 2008; Crisp *et al*., 2009; Donoghue & Edwards, 2014). Thus, niche conservation likely limits evolutionary transitions between biomes, especially between tropical and temperate biomes (Wiens & Donoghue, 2004; Crisp *et al*., 2009), which greatly differ in terms of environmental challenges (Folk *et al*., 2020). While rare, the gain of physiological adaptations to tolerate abiotic stress, such as freezing pressure, has been observed repeatedly across the angiosperm tree of life (Zanne *et al*., 2018; Folk *et al*., 2020). As a reflection of niche conservatism, it is thought that transitions from tropical to temperate biomes were facilitated by adaptations to seasonally dry tropical environments, because ancestral tropical lineages already possessed traits facilitating seasonal stressors such as drought complementary to the physiological traits needed for surviving freezes (Edwards *et al*., 2017; Folk *et al*., 2020). While such exaptations provided a means for tropical lineages to shift into temperate regions, these shifts were rare and it is more likely that lineage dispersals happened frequently between temperate regions, thereby contributing to the overall build-up of temperate biodiversity. Temperate biomes are also thought to differ in age across the globe, with old temperate biomes (e.g., sclerophyll biomes) tending to serve as a source for lineage dispersal to younger ones (e.g., arid, alpine, grassland) (Crisp *et al*., 2009; Donoghue & Edwards, 2014). Thus, work to date suggests that any of the three potential diversity drivers (*in-situ* diversification, dispersal, and time-for-speciation) may be responsible for centers of temperate diversity.

To understand the processes that underlie temperate centers of biodiversity, we examine historical eco-evolutionary dynamics through a suite of phylogenetic comparative methods with a focus on one angiosperm clade. We hypothesize (H1) that the colonization of temperate biomes happened independently and contemporaneously across lineages, due to global cooling and drying since the Oligocene leading to parallel origins of temperate environments on different continents. Second, we hypothesize (H2) that high global and regional species richness in our key clade is the result of high *in*-*situ* diversification rates, rather than high immigration rates or early colonization times. Specifically, we expect that temperate systems provided novel ecological opportunities or ‘adaptive zones’ (Simpson 1953) for increased diversification rates with limited immigration from tropical regions and relatively recent and similar colonization times across areas. This scenario down weights models assuming gradual diversity accumulation over time, since it explicitly suggests late origination and provides little time for temperate diversity to differentially increase among areas. Finally, we hypothesize (H3) that movement between temperate regions is relatively common, that is, there are strong source-sink dynamics that provide an engine for the unequal accumulation of lineages in today’s centers of diversity.

To test these hypotheses, we focus on the buckthorn family Rhamnaceae (Rosales), a clade with a nearly cosmopolitan distribution of ca. 1100 species within 63 genera (POWO, http://plantsoftheworldonline.org/). Rhamnaceae comprise predominantly woody shrubs with sclerophyllous leaves, exhibiting high diversity in fire-prone scrublands in temperate Mediterranean-type ecosystems, but also occurring in desert environments and temperate to tropical forests (Medan & Schirarend, 2004; Ladiges *et al*., 2005; Onstein *et al*., 2015; Onstein *et al*., 2016). We reconstructed and dated the most comprehensive Rhamnaceae phylogeny to date, with extensive sampling of genera and species (574 species in 58 genera, or ca. 52.2% and 92.0%, respectively) and genetic regions (89 low-copy nuclear loci). We then defined geographic- and climatic-based temperate biomes, and reconstructed the diversification history of Rhamnaceae across the temperate biome as a whole, within the most species-rich regions, and across local assemblages, in an effort to identify cross-scale macroevolutionary processes.

## Materials and Methods

### Plant sampling, sequencing, and data processing

We sampled 574 Rhamnaceae species (∼52.2% of the 1100 recognized species according to POWO) from 58 genera (of the 63 genera according to POWO), with representatives of all 11 tribes and 9 of 10 genera unassigned to any tribe (Richardson *et al*., 2000a; Richardson *et al*., 2000b; Hauenschild *et al*., 2016). Three species of Elaeagnaceae and one species each of Barbeyaceae and Dirachmaceae (Rosales) were included as outgroups (Li *et al*., 2021).

Leaf material was collected from the following herbaria: A, AD, BRI, CAS, F, KUN, MEL, MO, NY, OS, PERTH, TEX, and US as well as from the field (Dataset S1), and total DNA was extracted using a modified CTAB method following the protocol described in Folk *et al*. (2021). Target enrichment probes (Folk *et al*., 2021; Fu *et al*., 2022) were used to capture 100 low-copy nucleotide genes, and hybridization enrichment sequencing (Hyb-seq) was conducted by Rapid Genomics (Gainesville, Florida, USA). Raw sequenced reads were cleaned and filtered as follows: Illumina adapter sequence artifacts were trimmed, low-quality reads were discarded, and low-quality read ends were trimmed using TRIMMOMATIC v0.32 (Bolger *et al*., 2014).

Assembly of the processed nuclear reads was performed using HybPiper v1.2 (Johnson *et al*., 2016), a reference-based assembler, using the 100 protein sequences from *Arabidopsis thaliana* used for probe design as the reference. Reads were mapped to each reference using BLASTX v2.7.1 (Camacho *et al*., 2009), each gene was assembled *de novo* using SPAdes v3.12.0 (Bankevich *et al*., 2012) and coding sequences were extracted using Exonerate v2.4.0 (Slater & Birney, 2005). Each gene missing > 75% of the sampled species was excluded. As a result, 89 loci were kept for further analysis.

### Phylogenetic analyses and divergence time estimation

The sequences of each targeted gene region were initially aligned using MAFFT (Katoh & Standley, 2013) using default settings. To reduce errors in our alignments (i.e., gap-heavy and ambiguously aligned sites), we cleaned the original alignment of each gene using ‘pxclsq’ in phyx (Brown *et al*., 2017), removing alignment columns with < 30% occupancy. The cleaned alignments were concatenated into a supermatrix using the ‘pxcat’ function in phyx. This supermatrix (Dataset S2) was then used to infer phylogenetic relationships in Rhamnaceae using the GTR-GAMMA model with 1,000 bootstrap replicates in RAxML v8.2.11 (Stamatakis, 2014). In addition, a coalescent species tree was inferred from the 89 best maximum likelihood (ML) single-gene trees, which were built in RAxML v8.2.11 using the GTR-GAMMA model with 200 bootstraps (BS), using ASTRAL-III v5.6.3 (Zhang *et al*., 2018). Branches with < 10% BS in each gene tree were collapsed using Newick utilities (Junier & Zdobnov, 2010).

The concatenated supermatrix and the corresponding ML tree were used for dating analysis. We estimated divergence times using the penalized likelihood method implemented in treePL (Smith & O’Meara, 2012). Six fossils and a secondary calibration were used to calibrate node ages (more details in Method S1). We used an optimal smoothing parameter determined by the “random subsample and replicate” cross-validation method to accommodate rate heterogeneity. To assess uncertainty in age estimates, we estimated confidence intervals on inferred ages by dating all 100 ML bootstrap trees from the concatenated dataset (Maurin, 2020). Results from the dating of the bootstrapped trees were then summarized and visualized on the concatenated ML tree using TreeAnnotator v2.6.3 (Bouckaert *et al*., 2014).

### Collection of species occurrence data and environmental variables

To illustrate the distribution and species richness of Rhamnaceae across temperate biomes, we collected occurrence data of all Rhamnaceae species from the global biodiversity information facility (GBIF, https://www.gbif.org/). The World Checklist of Vascular Plants (WCVP, http://wcvp.science.kew.org/) was used to standardize species with accepted names, and the infraspecific taxa and exotic and hybrid species were excluded. POWO was used to provide the native status for each species. These occurrence datasets were carefully assessed, and records that lacked geographic coordinates, occurred in the oceans and were duplicates were removed by customized R scripts. Additionally, cultivation records were removed manually. Finally, a total of 291,041 unique distribution records from 1022 Rhamnaceae species were used in subsequent steps (Dataset S3).

To classify the temperate biomes and associated environmental conditions, we collected 35 environmental variables, including 19 bio-climatic variables at 2.5m resolution (https://www.worldclim.org), two topographical layers (https://lta.cr.usgs.gov/GTOPO30), eight soil layers (averaged in QGIS across layers at 5-, 15-, 30- and 60-cm sampling depths; https://soilgrids.org/), and six land cover classes (https://www.earthenv.org/landcover). All environmental variables were extracted as mean values per 1° × 1° grid cells using the zonal statistics in the QGIS v3.2 (QGIS Development Team, 2018). Mean values of the variables for each grid cell were used in the subsequent environmental niche analysis (Dataset S4).

Global patterns of Rhamnaceae species diversity, temperate biomes, and biodiversity ‘hotspots’

To identify the global pattern of Rhamnaceae species richness, we matched the global filtered distribution dataset to the grid cells of 1° × 1° based on presence/absence of species within each grid cell (Dataset S4). We then calculated the species richness of each grid cell using the R package ‘phyloregion’ (Daru *et al*., 2020).

To classify grid cells as occurring in temperate or tropical (non-temperate) biomes, we used geographic and climatic definitions for the tropics (Peel *et al*., 2007; Owens *et al*., 2017). According to the geographic definition, we classified grid cells falling outside the range between 23.4°N and 23.4°S (the Tropics of Cancer and Capricorn) as temperate, while classified those within this range as tropical. According to the Köppen-Geiger climatic definition (Peel *et al*., 2007), we classified grid cells with a year-round monthly mean temperature < 18 °C as temperate and classified those with a temperature ≥ 18 °C as tropical (Dataset S4). In addition, we used these two definitions to classify each Rhamnaceae species as occurring in temperate and/or tropical biomes, while considering occurrence frequency, i.e., species with >70% of their occurrences in the temperate region were classified as temperate; species with >70% of their occurrences in the tropics were classified as tropical; otherwise, species were classified as both (temperate + tropical) (Dataset S5). We associated all grid cells and species occurrence points with average monthly annual mean temperature data using bio1 from WorldClim (Fick & Hijmans, 2017).

We conducted an optimization analysis using the Gi* local statistic (Getis & Ord, 1996) implemented in QGIS v3.2 with the Hotspot Analysis plugin (Oxoli *et al*., 2017) to identify regions particularly species-rich, hereafter Rhamnaceae ‘hotspots’. This method calculated the local spatial autocorrelation to cluster grid cells of high species richness. We identified each hotspot as a region consisting of adjacent grid cells with Z scores ≥ 1.65 (Suissa *et al*., 2021), enforcing at least 30 grid cells per hotspot to ensure that each hotspot has a sufficient sample size and spatial extent. Once hotspots were identified, we defined non-hotspot regions as all remaining grid cells with at least one Rhamnaceae species (Dataset S4). Finally, we assessed whether the same hotspots were identified when we only included Rhamnaceae species included in the phylogenetic tree. Therefore, we calculated Spearman’s correlation between grid-based species richness when including all species distribution data (n = 1022 species) to species richness when including only phylogenetically sampled species (n = 574 species) (Spearman’s ρ = 0.93 in our dataset).

### Environmental niche of temperate biomes and biodiversity hotspots

To assess whether high Rhamnaceae species richness was linked to environmental conditions typical for temperate biomes, and to assess whether hotspots shared such environmental space, we performed Principal Coordinate Analyses (PCoA). All environmental variables (i.e., climate, topography, soil and landcover) were log-transformed to improve homoscedasticity, and we scaled them between 0 and 1 to impose equal variable weight. PCoA was computed to describe the composition of environmental space of each hotspot region, using Euclidean dissimilarities calculated in the R packages ‘vegan’ (Dixon, 2003) and ‘ape’ (Paradis *et al*., 2004). The volume of the convex hull composed by two independent axes (PCoA1 and PCoA2, collectively capturing 59.6% of the environmental variation) was then used to measure the overlap in climatic hypervolume space of temperate biomes and for each Rhamnaceae hotspot region.

### Inference of biogeographic history of biodiversity hotspots

To assess the independent and contemporaneous colonization of temperate regions (H1) and subsequent diversity accumulation through time (H2), we traced the biogeographic history of Rhamnaceae lineages across the temperate hotspots. Ancestral ranges were inferred by implementing a series of biogeographic models in the R package BioGeoBEARS v1.1.2 (Matzke, 2013). Outgroups were excluded from the time-calibrated phylogenetic tree. We used the corrected Akaike Information Criterion (AICc) to compare the fit of three biogeographical models: DEC, DIVALIKE, BAYAREALIKE and versions of these three models that allowed for founder event speciation (+J) (Matzke, 2014), i.e., DEC+J, DIVALIKE+J, and BAYAREALIKE+J. We defined eight areas, following our identification of Rhamnaceae hotspot regions: California, Mexico to Central America, Mediterranean Basin, Southern China, South African Cape, Southwest Australia, Southeast Australia and non-hotspot. Species were assigned to one or more biogeographic areas based on occurrence frequency within these regions, with presence only being assigned if species with > 10% of its occurrences fell within a particular biogeographic area. The maximum number of areas was set to three, to reflect distributions of extant taxa.

We performed 100 biogeographical stochastic mappings (BSMs) (Dupin *et al*., 2016) with the parameter rate estimates of the best-fitting model from the BioGeoBEARS analysis. The BSMs produced a probabilistic sample of the chronology of anagenetic and cladogenetic events, which allowed us to count and date anagenetic dispersals, extinctions, cladogenic range expansions (i.e., sympatry), and founder events. In addition, we binned these events into 0.5-Ma periods and calculated the 95% confidence interval of the number of lineages occupying each region at each time step from 100 BSMs using the R package ‘ltstR’ (Skeels, 2019). Then, we extracted median values from the 95% confidence intervals to calculate the relative proportion of lineages in each hotspot and non-hotspot at each time bin.

### Diversification and dispersal rates of biomes and biodiversity hotspots

To assess whether high *in*-*situ* diversification rates explained high Rhamnaceae species richness in temperate biomes and hotspots (H2) and whether the dispersal of lineages between temperate regions was relatively common (H3), we estimated speciation, extinction, and dispersal rates with the Geographic State Speciation and Extinction (GeoSSE) model (Goldberg *et al*., 2011) implemented in the R package ‘diversitree’ (FitzJohn, 2012). To this end, we used the classification of species as assigned above as ‘temperate’, ‘tropical’, or ‘both’. Furthermore, we also fitted GeoSSE models in which we classified species for each hotspot region (California, Mexico to Central America, Mediterranean Basin, Southern China, Cape, Southwest Australia, Southeast Australia) against species occurring elsewhere, i.e., seven analyses in total. ‘Elsewhere’ therefore includes all other recognized hotspots except the specific hotspot as well as Rhamnaceae occurrences in ‘non-hotspot’ regions. Each species was assigned to a hotspot based on its occurrence frequency (i.e., species with >90% of occurrences within a specific hotspot were assigned to that hotspot). Species with >90% of their occurrences outside any recognized hotspot were classified as occurring ‘Elsewhere’; finally, species were classified as widespread if they occurred both in specific hotspots and elsewhere. In addition, we ran another analysis in which we contrasted species in any hotspot region to species in non-hotspot areas.

All diversification analyses were conducted on the time-calibrated phylogenetic tree without outgroups. We compared the fit of eight models that allowed speciation, extinction, and dispersal rates to vary between temperate and tropical biomes or between lineages in hotspots vs. elsewhere and selected the best-fitting model as assessed by a likelihood ratio test. The best-fit model was then used in a subsequent Bayesian MCMC run for 5000 generations (ESS > 200), using ML rate estimates as starting points and an exponential prior whose distribution was in relation to the overall diversification rate, estimated using the Kendall–Moran estimate for net diversification rate (Kendall, 1949; Moran, 1950). Because species in the Mediterranean Basin hotspot and the Southwest Australian hotspot consisted of <10% of the total species richness of Rhamnaceae, we were not able to infer rates for these hotspots individually due to sample size constraints (Davis *et al*., 2013), but they were included in the analysis in which all hotspots were combined and compared to diversification and dispersal rates elsewhere.

### Phylogenetic assemblage structure of biodiversity hotspots

Another signature of high diversification rates can be captured by phylogenetic clustering – i.e., species within a local assemblage are phylogenetically more closely related than expected by chance, suggesting *in*-*situ* diversification from a common ancestor. We therefore calculated the Net Relatedness Index (NRI) (Webb *et al*., 2002) for each grid cell that contained more than one Rhamnaceae species (n = 3266). NRI measures how mean phylogenetic distance (MPD) between all species pairs in the grid cell deviates from random. The calculation of NRI and associated 999 randomization tests were conducted with the R package ‘picante’ (Kembel *et al*., 2010), and values were compared between assemblages in temperate and tropical biomes and hotspots.

### Sensitivity of macroevolutionary processes to hotspot definitions

To test whether diversification rates, dispersal rates and colonization time inferences were robust to the size of hotspot delimitations from the Gi* local statistic, we used a distance of 1 degree as a buffer distance around each hotspot (hereafter buffer-hotspots), and repeated the BioGeoBEARS, BSMs, and GeoSSE, analyses with this new classification. All analyses above were carried out in R (R Core Team, 2022), unless mentioned otherwise.

## Results

### Phylogenetic analyses and divergence time estimation

After assembling and filtering, 89 low-copy nuclear genes were obtained for 574 Rhamnaceae species and five outgroup species. The concatenated data matrix was 93,936 bp in length. The three main Rhamnaceae groups, i.e., the rhamnoid group, the ziziphoid group, and the clade containing several Rhamnaceae taxa of few genera (*Bathiorhamnus*, *Doerpfeldia*, *Sarcomphalus*, *Ziziphus*) were fully supported in the phylogeny (BS = 100%; LPP = 1) using both concatenation and coalescent methods (Figs. 1a, S1, S2). The concatenated ML tree and coalescent ASTRAL tree were largely congruent, with most of the deeper nodes and branches resolved and strongly supported, except for a few nodes within the ziziphoid clade (Figs. S1, S2). These showed conflicts and obtained relatively low support. Rhamnaceae were estimated to have originated at around 113.54 Ma (113.33–113.60 Ma; 95% highest posterior density) in the Cretaceous based on the concatenated ML tree, and details for the age estimates of clades are presented in Fig. S3.

**Fig. 1.**
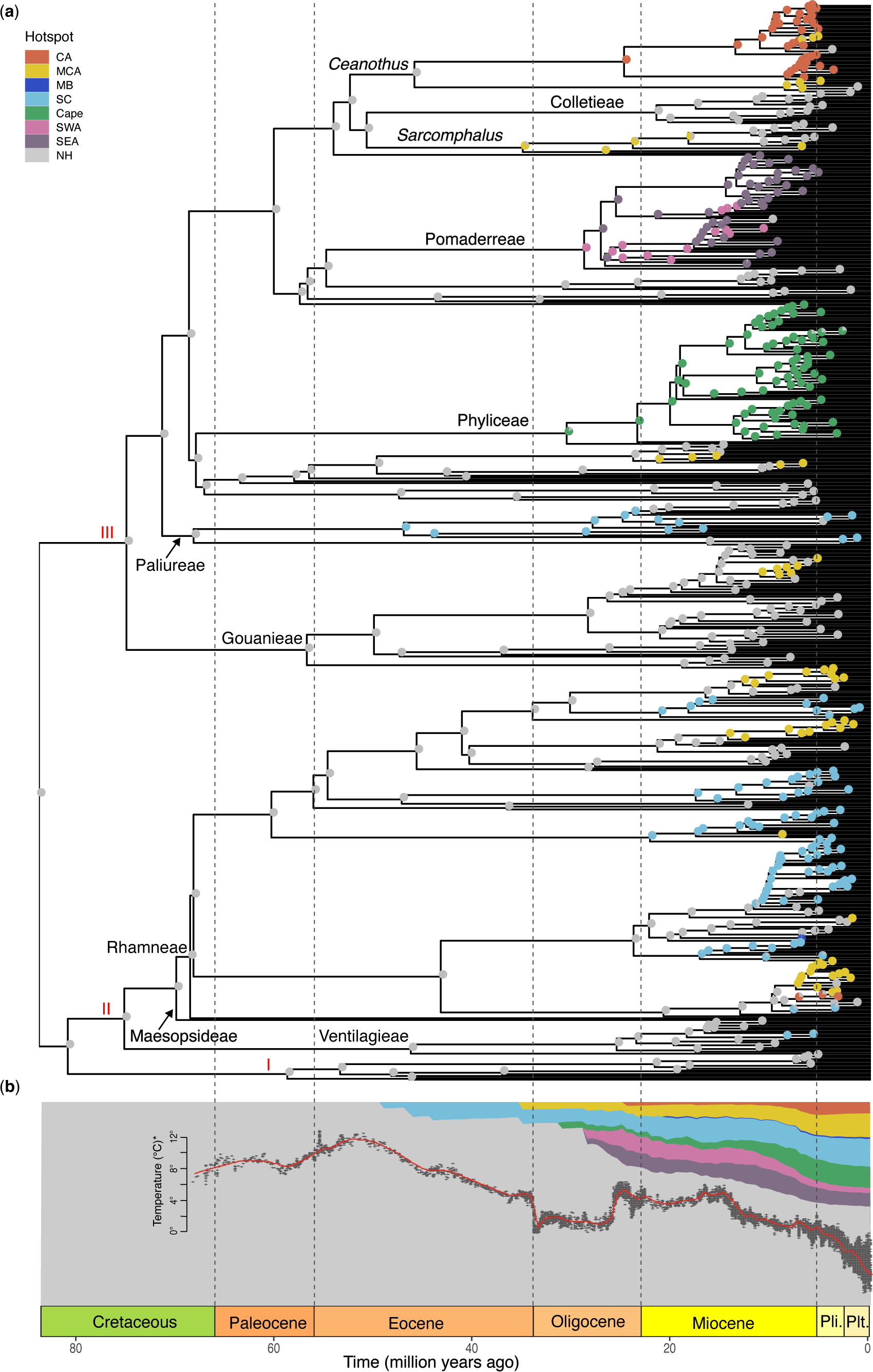
Ancestral range estimates and time-for-speciation of Rhamnaceae lineages in biodiversity hotspots. (a) Divergence times estimated using treePL based on the concatenated supermatrix of 89 low-copy loci, and the ancestral areas reconstructed by BioGeoBEARS using the BAYAREALIKE + J model. I, the group containing genera (*Bathiorhamnus*, *Doerpfeldia*, *Sarcomphalus*, *Ziziphus*); II, the rhamnoid group; III, the ziziphoid group. CA, California; MCA, Mexico to Central America; MB, Mediterranean Basin; SC, Southern China; SWE, Southwest Australia; SEA, Southeast Australia; NH, non-hotspot. (b) Colonisation and time-for-speciation of Rhamnaceae lineages in biodiversity hotspots. The relative proportion of lineages in each hotspot region through time, estimated from the biogeographical stochastic mappings (BSMs) analysis using a sliding window and median value from the 95% confidence interval of the number of lineages occupying each region at each time bin. Red line is global temperature curve inferred from Zachos et al. (2001) over the past 65 million years. Pli., Pliocene; Plt., Pleistocene.

### Global patterns of species diversity, temperate biomes and biodiversity hotspots in Rhamnaceae

Based on the global species richness map (Fig. 2a), Rhamnaceae species diversity was markedly higher in temperate (mid-latitude) than in tropical (low-latitude) regions (Fig. 2b), resulting in a bimodal latitudinal diversity gradient. Rhamnaceae assemblages and species were predominantly temperate based on geography (60.85% assemblages and 61.06% species) but not based on temperature (50.45% assemblages and 49.51% species; Datasets S4, S5). Some species were scored as belonging to both temperate and tropical areas (4.30% and 12.16% based on geographic and climatic definitions, separately), so tropical percentages were lower than temperate for both definitions. We identified seven distinct regions as hotspots with exceptionally higher species richness than non-hotspots (Z scores ≥ 1.65; Figs. 2a, S4; Dataset S4), i.e., California (n = 60 [grid cells]), Mexico to Central America (n = 224), Mediterranean Basin (n = 57), Southern China (n = 221), Cape (n = 33), Southwest Australia (n = 51), and Southeast Australia (n = 126). Grid cells outside the seven hotspots, defined as non-hotspots, were more numerous (n = 5254).

**Fig. 2.**
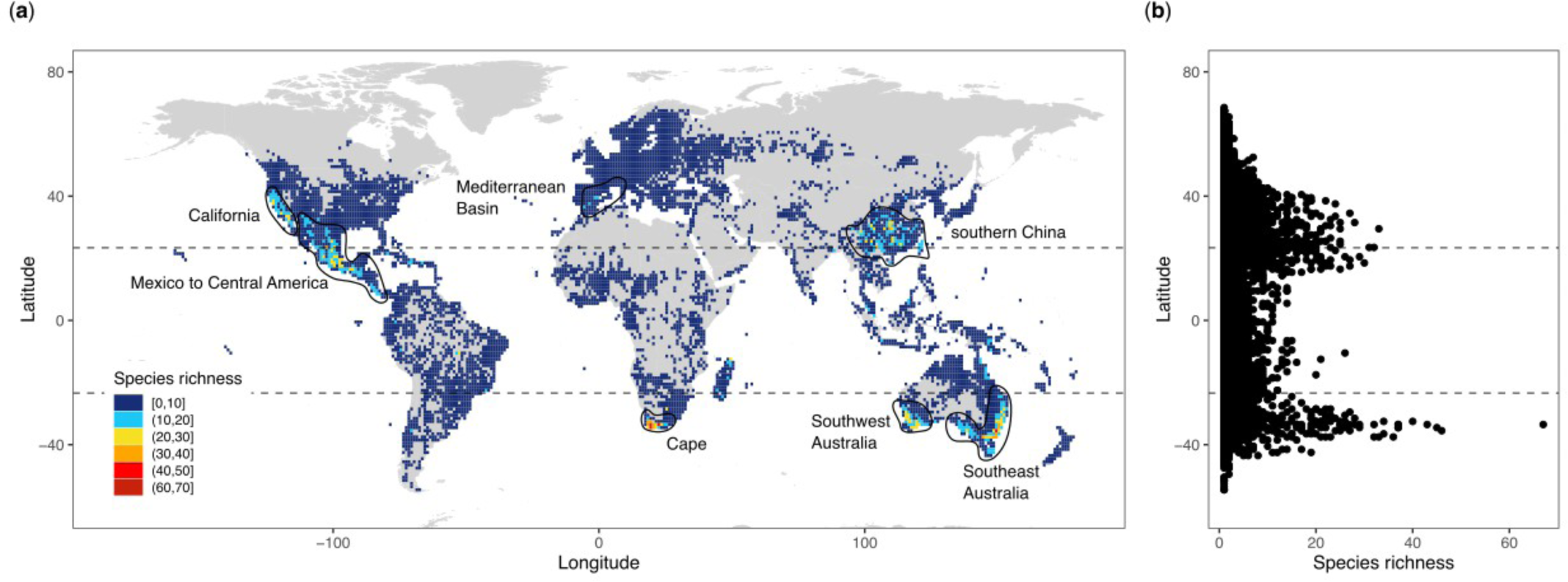
The global distribution of Rhamnaceae species richness across assemblages in the temperate realm. (a) Rhamnaceae species richness estimated from 291,041 unique distribution records of 1022 species. Colored squares indicate estimated species richness in grid cells of 1° × 1°. Regions containing hotspots of Rhamnaceae diversity outlined with solid lines. (b) Species richness across latitude. Each point represents the species richness within a 1° × 1° grid cell plotted against the grid cell’s latitudinal centroid.

### Similarities in environmental features across temperate Rhamnaceae hotspots

We quantified the hypervolume of the environmental space as defined by PCoA1 (explaining 38.9% of the variance) and PCoA2 (20.7%) in temperate biomes and each hotspot (Fig. 3). The PCoA showed that environmental space of all hotspots was nested within the temperate biome environmental space according to both geographic and climatic definitions (Figs. 3, S5a), with the exception of parts of Mexico to Central America. Furthermore, the PCoA showed substantial overlap (on average 14.8%–56.4% per hotspot; Table S1) among the seven hotspots in environmental hypervolume space. A biplot showed that temperature variables (e.g., isothermality and annual mean temperature) along the PCoA1 axis and soil variables (e.g., soil organic carbon and soil pH) along the PCoA2 axis were the most important predictors of the environmental hypervolume (Fig. S5b). The biplot also showed tropical climatic features (right side of the biplot) in Mexico to Central America, nutrient-rich soils in the Southeast Australia hotspot (top side of the biplot), and colder temperatures in the Southern China hotspot (left side of the biplot), that distinguish environments in these hotspots from the other hotspots.

**Fig. 3.**
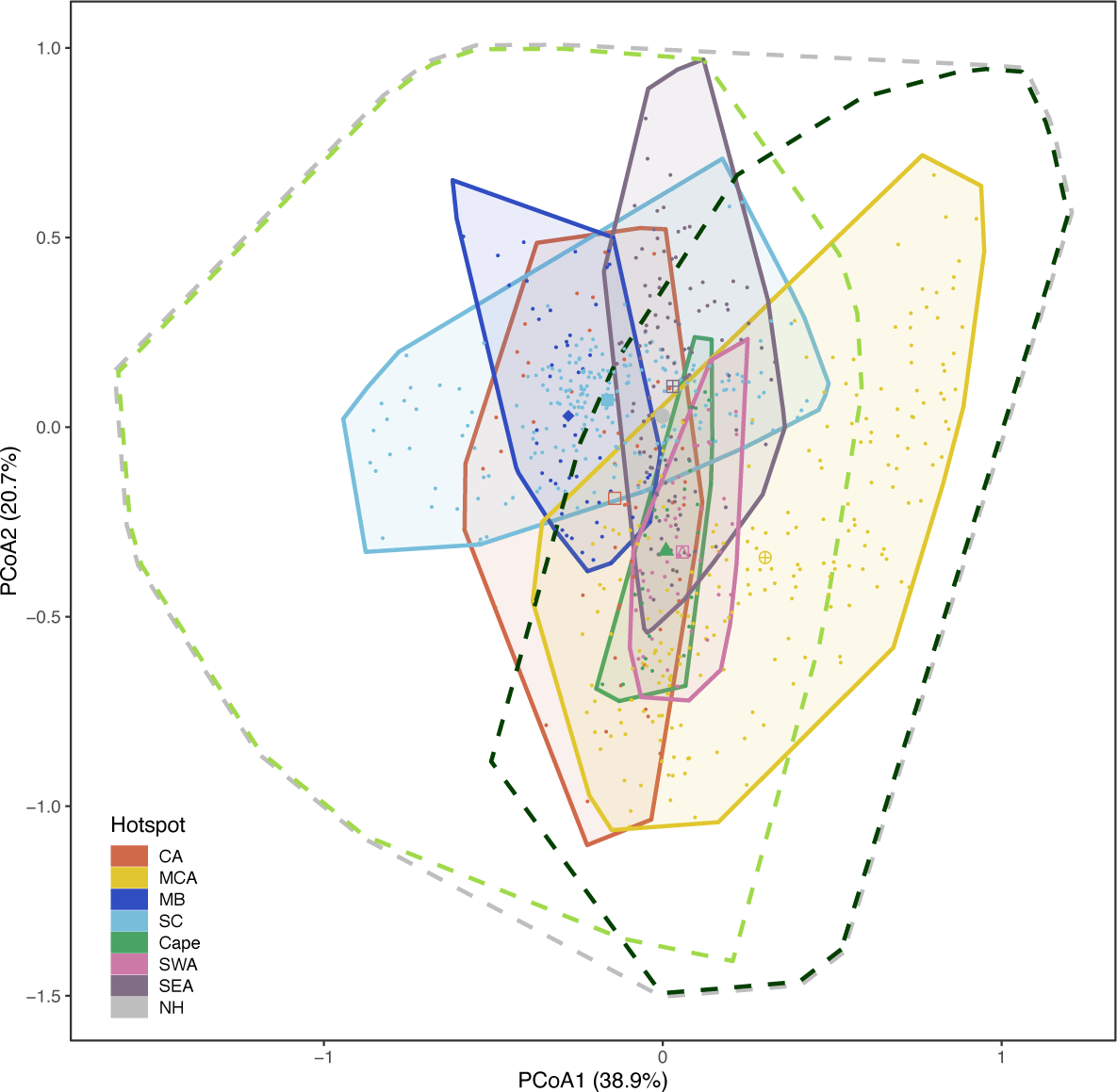
The occupancy of hypervolume environmental space in Rhamnaceae temperate and tropical biomes. Colored polygons are two-dimensional environmental hypervolumes based on 772 assemblages (grid cells) in seven hotspots using principal coordinates analysis (PCoA). The light green, dark green, and gray dashed polygons are the environmental hypervolumes of respectively, temperate, tropical (according to the geographic definition), and non-hotspot regions. Colored points represent grid cells. Colored shapes are hypervolume centroids. CA, California; MCA, Mexico to Central America; MB, Mediterranean Basin; SC, Southern China; SWE, Southwest Australia; SEA, Southeast Australia.

### Historical biogeography of hotspots

The BAYAREALIKE + J model was selected in the ancestral biogeographic reconstruction of Rhamnaceae hotspots based on the AICc (Table S2; Fig. S6). By comparing the relative accumulation of lineages through time in each hotspot (Fig. 1b), we detected the earliest colonization of the temperate region at ca. 49.0 Ma (46.5–83.5 Ma; 95% confidence interval of age of lineages occupying each region from 100 BSMs), i.e., with the colonization of Southern China during the Eocene. Subsequently, lineages independently colonized Mexico to Central America at ca. 35 Ma (29.5–83.5 Ma), the Cape at ca. 31.0 Ma (23.0–83.5 Ma), Southwest Australia at ca. 28.5 Ma (28.5–83.5 Ma), Southeast Australia at ca. 28.5 Ma (27.0–57 Ma), and California at ca. 24.5 Ma (24.5–62.0 Ma). Thus, Rhamnaceae achieved their modern distribution through a series of largely contemporaneous dispersals during the Oligocene, consistent with H1. Lineages colonized the Mediterranean Basin somewhat later during the early Miocene, ca. 20.5 Ma (7.0–83.5 Ma). We found that the proportion of Rhamnaceae lineage diversity in the hotspot regions compared to elsewhere continued to increase rapidly and in parallel across hotspots from the Miocene onwards. The BAYAREALIKE model was selected in the biogeographic analysis of buffer-hotspots (Table S2), but ancestral biogeographic reconstruction, colonization times, and lineage accumulation curves in buffer-hotspots were similar overall to those in the main hotspot analysis (Figs. S6–S8).

### Macroevolutionary rates of diversity in hotspots

The best-fitting GeoSSE models indicated distinct diversification and/or dispersal rates for temperate vs. tropical biomes, and for each hotspot vs. elsewhere, based on likelihood ratio tests (Table S3). Lineages in temperate biomes showed higher speciation and net diversification rates than lineages in tropical biomes, a result that was robust to geographic (Figs. 4a–b) and climatic (Figs. S9a–b) definitions of temperate biomes. Geographic and climatic definitions did differ in extinction rates, with lower extinction rates in geographically temperate areas but equal extinction rates in climatically temperate areas. Furthermore, lineages occurring in hotspots showed higher net diversification rates than lineages occurring elsewhere, except Southern China, thus mostly consistent with H2, but speciation and extinction rates differed among the five hotspots (Figs. 4e–g). Specifically, California, Cape, and Southeast Australia showed higher speciation rates, resulting in higher net diversification rates for lineages in these hotspots than lineages in Mexico to Central America and Southern China. For Mexico to Central America, the lowest speciation rates resulted in relatively low net diversification rates. For Southern China, the relatively low speciation rates and the highest extinction rates resulted in the lowest net diversification rates across hotspots, similar to diversification rates in non-hotspots. Rates of speciation, extinction, and net diversification in buffer-hotspots were qualitatively similar to those in hotspots (Figs. 4e–g, S9d–f).

**Fig. 4.**
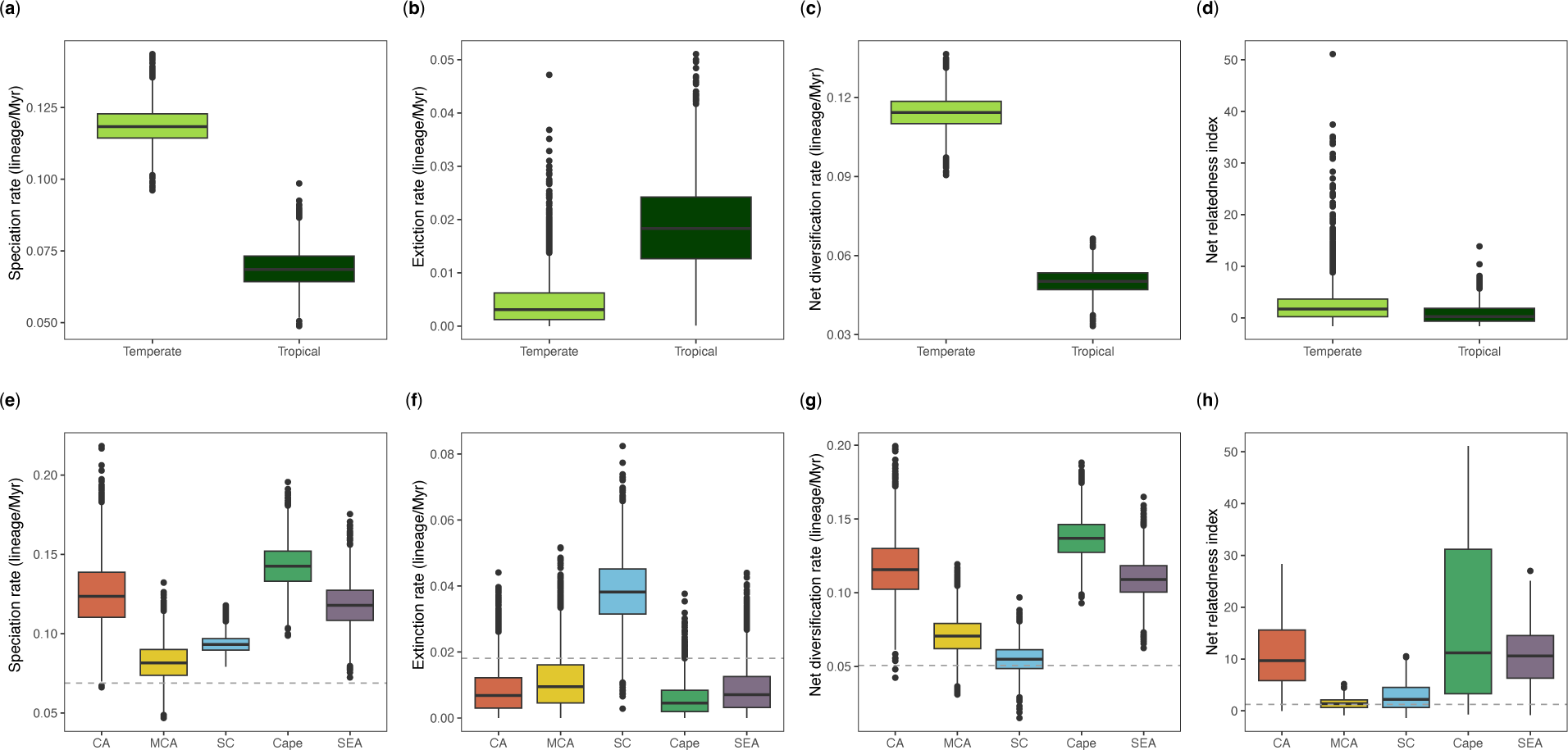
Diversification rates and net relatedness index across biomes in Rhamnaceae. Estimated parameter distributions in temperate vs. tropical biomes (according to the geographic definition) as well as of Rhamnaceae lineages evolving in five hotspots. The Mediterranean Basin and Southwest Australia were excluded because sampling of species in these two hotspots was less than 10% compared to elsewhere. Boxplots showing (a) speciation rates (lineage/Myr), (b) extinction rates (lineage/Myr), (c) net diversification rates (lineage/Myr), and (d) net relatedness index (NRI) of temperate and tropical biomes; (e) speciation rates (lineage/Myr), (f) extinction rates (lineage/Myr), (g) net diversification rates (lineage/Myr), and (h) net relatedness index (NRI) of the five hotspots. Boxes in figures (a-c, e-g) are colored by region and represent parameter distributions from the Bayesian MCMC using the best-fitting GeoSSE model on the time-calibrated Rhamnaceae phylogenetic tree. Dashed lines shown in figures (e-h) are averaged parameter values of non-hotspot lineages. CA, California; MCA, Mexico to Central America; SC, Southern China; SEA, Southeast Australia.

GeoSSE results showed equal migration and immigration (i.e., dispersal) rates between temperate and tropical regions according to the geographical definition (Fig. 5). However, the climatic definition of regions resulted in estimated dispersal rates from temperate biomes to tropical biomes, and from hotspots to elsewhere, as significantly higher than *vice versa* in all five hotspots (Figs. 5, S10), partly consistent with H3. Among hotspots, we found that dispersal rates out of Mexico to Central America and Southern China were comparatively high, while dispersal rates out of the Cape and Southeast Australia hotspots were relatively low. Rates of dispersal in buffer-hotspots were qualitatively similar to those in hotspots (Fig. S10).

**Fig. 5.**
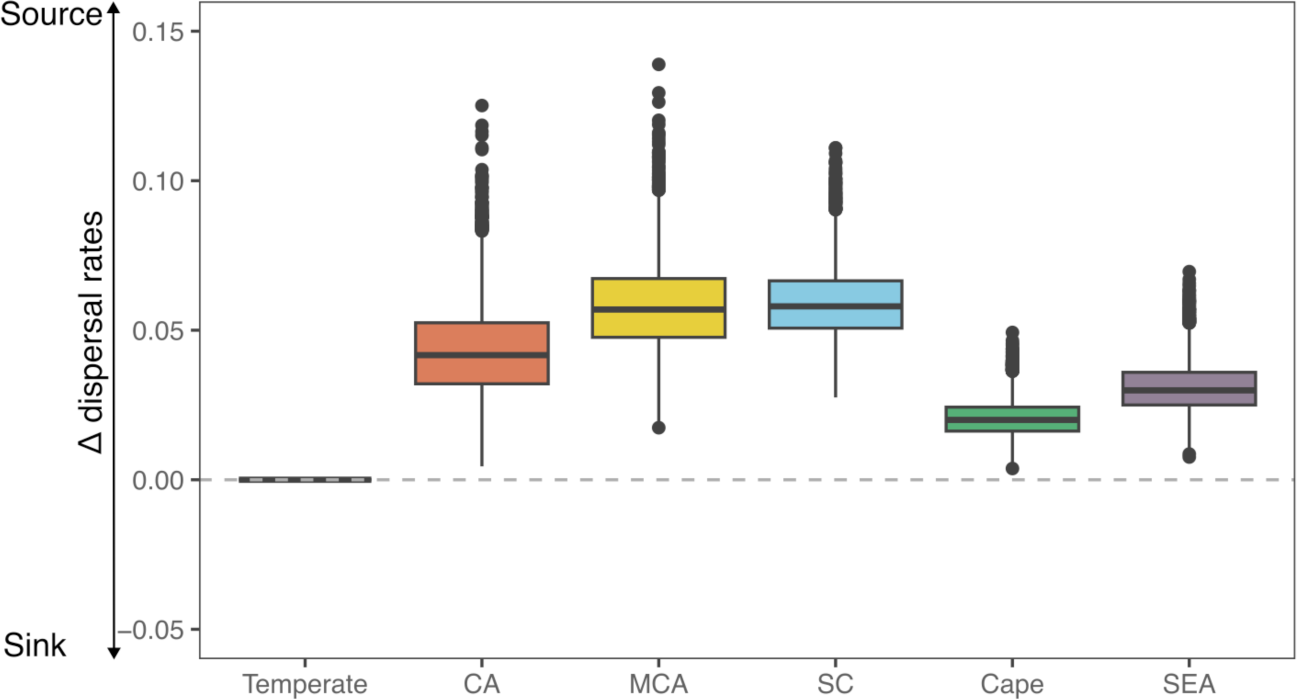
Source and sink dynamics of Rhamnaceae in temperate regions. Boxes are colored according to temperate biomes (according to the geographic definition) and hotspots and represent parameter distributions (dispersal rates out of a region minus dispersal rates into that region) from the Bayesian MCMC from the best-fitting GeoSSE model on the time-calibrated Rhamnaceae phylogenetic tree. CA, California; MCA, Mexico to Central America; SC, Southern China; SEA, Southeast Australia.

### Phylogenetic clustering of lineages in hotspots

NRI showed significant phylogenetic clustering in temperate compared to tropical biomes, as well as within assemblages in California, Cape, and Southeast Australia (Figs. 4d, g, S13), suggesting that these assemblages are composed of closely related species, thus consistent with high, primarily *in*-*situ* diversification rates in these hotspots (Fig. 4). The NRI comparison among the seven hotspots indicated that both Mexico to Central America and Southern China had an overall random/overdispersed structure with relatively low NRI compared with those of the other hotspots, suggesting that assemblages are instead composed of distantly related species that may reflect dispersal.

## Discussion

We dissected the intricate relationships between environment, diversification, and dispersal to understand high biodiversity in temperate biomes, using Rhamnaceae as a model system. We identified seven hotspots particularly high in Rhamnaceae species richness: California, Mexico to Central America, Mediterranean Basin, Southern China, Cape, Southwest Australia, and Southeast Australia (Figs. 2a, S4). With some differences among particular hotpots, our results overall point to high *in*-*situ* diversification rates (Figs. 4, S9), rather than high immigration rates or accumulative time (Figs. 1b, 5, S9, S10) as the primary process behind the high diversity of Rhamnaceae in temperate biomes.

We show that all Rhamnaceae hotspots overlap in current environmental space, particularly in the climatic facets most important for defining the unique stressors of temperate – and especially Mediterranean – ecosystems (Figs. 3, S5). Notably, California, the Mediterranean Basin, Cape, and Southwest Australia all share a Mediterranean climate with dry, hot summers, cooler, wet winters, and fire-prone and woody shrubland vegetation (Donoghue & Edwards, 2014; Rundel *et al*., 2016). This overlap in environment among Rhamnaceae hotspots, which comprise phylogenetically disparate species clusters and independent colonizations from tropical ancestors during the Oligocene (Fig. 1b), suggests a possible role of evolutionary convergence in building diversity across hotspots. As well, Mediterranean environments may have selected for lineages possessing functional traits that facilitate survival under seasonal drought and low-nutrient soils, e.g., small, evergreen, sclerophyllous leaves of Rhamnaceae species facilitate survival on seasonally dry and old, climatically buffered, infertile landscapes (OCBILs per Hopper, 2009; Onstein *et al*., 2016; Ackerly & Onstein, 2018).

Rhamnaceae colonizations of several geographically separated temperate regions show a striking similarity in timing (i.e., 35–24.5 Ma), but lineage diversity increases in these biomes later, particularly so from 23 Ma onwards (Figs. 1b, S8). We argue that diversity within the Mediterranean temperate hotspots has resulted from a set of *in*-*situ* – possibly nested – radiations, where a combination of global climate change and pre-adapted traits (e.g., sclerophyllous leaves) facilitated parallel and rapid diversification across seasonal and xeric climatic regions (Onstein *et al*., 2016; Rundel *et al*., 2016). This is consistent with the global expansion of temperate biomes from the Oligocene onwards (34 Ma) due to strong climate cooling events and drying, providing ecological opportunities for temperate biodiversity to originate and expand. Similar patterns of increased diversification coinciding with the onset of colder and drier climates in the Miocene have been found in some angiosperm lineages, such as Poaceae, Asteraceae, Saxifragales, rosids, and orchids (Folk *et al*., 2019; Soltis *et al*., 2019; Sun *et al*., 2020; Palazzesi *et al*., 2022; Thompson *et al*., 2023a).

Lineages in California, Cape, and Southeast Australia showed high speciation and low extinction rates and strong phylogenetic clustering of assemblages (Figs. 4d–g). Our results are consistent with the many evolutionary radiations characterized in Rhamnaceae in previous work, such as *Ceanothus* (ca. 60 species) in Californian chaparral (Burge et al., 2011), Phyliceae (ca. 150 species) in Cape fynbos (Linder, 2003), and Pomaderreae (ca. 240 species) in Australian shrublands (Kellermann, 2020; Nge et al., 2021). Furthermore, Rhamnaceae share this signature of high diversification with other lineages in these regions, such as *Arctostaphylos* (Ericaceae) in California (Stebbins & Major, 1965), *Moraea* (Iridaceae) in the Cape (Goldblatt et al., 2002), and *Acacia* (Caesalpinioideae) in Australia (Renner et al., 2020). Although Mexico to Central America showed phylogenetic overdispersion, and it harbors distinct Rhamnaceae lineages (*Ceanothus*, *Colubrina*, *Sarcomphalus*, and Rhamneae), these lineages showed overall high net diversification rates compared to Rhamnaceae lineages evolving elsewhere (Figs. 4d–g). It is likely that the phylogenetic overdispersion here may be related to high mixed endemism and high diversity of Mexico to Central America, as this region is at a crossroads between temperate and tropical regions and also highly heterogeneous environments because of its topographic complexity - a unique combination (Sosa *et al*., 2018).

The exception to the high *in*-*situ* diversification of Rhamnaceae in hotspots is Southern China. Here, lineages showed relatively low speciation rates and high extinction rates, resulting in much lower net diversification rates than lineages in any of the other hotspots, similar to rates in non-hotspot areas (Figs. 4d–g). Furthermore, assemblages in Southern China showed phylogenetic overdispersion (Fig. 4g), comprising distinct Rhamnaceae lineages (e.g., *Rhamnus*, *Berchemia*, *Rhamnella*, and *Ziziphus*), and the region was colonized much earlier than the other regions (Figs. 1b, S8). High species richness in Southern China may therefore be explained by the time-for-speciation hypothesis, with gradual accumulation of species diversity since the Eocene (Yan *et al*., 2018). It has been suggested that extinction over time maybe the reason for phylogenetic overdispersion in Southern China (Zhang *et al*., 2022). Indeed, Southern China is a center of paleo- and mixed paleo/neo-endemism of woody plants (Wang *et al*., 2022) and acts as refugia (i.e., paleo-endemism) for rare Chinese angiosperms, characterized by magnoliids and other ancient angiosperm lineages that survived extinction events and persist in a much narrower area than previously or places of more recent diversification (i.e., neo-endemism) (Lu *et al*., 2018; Wang *et al*., 2022; Zhang *et al*., 2022). Orogenic movements, annual temperature and annual precipitation may have experienced little change in mountainous areas of this region since the Cretaceous (Lu *et al*., 2018), and long-term climate stability may have provided the opportunity for some but not all lineages to persist in this refugial landscape. Thus, our results for Rhamnaceae provide macroevolutionary evidence that is consistent with Southern China, but not other hotspots, providing the long-term stability to accumulate diversity over geological time.

Our GeoSSE analysis showed that dispersal rates out of temperate biomes were equal or higher than out of tropical biomes overall, and dispersal rates into hotspots were significantly lower than dispersal rates out of hotspots (Figs. 5, S10) (also see Onstein *et al*., 2015), suggesting that while hotspots differed in dispersal rates, it is unlikely that high immigration rates have influenced the globally high Rhamnaceae species richness in temperate biomes. In addition, our results suggest that Rhamnaceae temperate hotspots may have acted as sources for recruitment of species in neighboring areas, making them diversity ‘source’ rather than ‘sink’ regions. The creation of such a source may be linked to the high *in*-*situ* diversification rates in these regions, resulting in high numbers of lineages and species. Our results suggest that these dispersals may have been to both temperate and tropical biomes (Figs. 5, S10–12). Thus, reversals to more tropical systems (e.g., seasonally dry tropical biome) from temperate regions are not uncommon in Rhamnaceae, probably because the temperate biomes they dominate are characterized by (seasonal) drought, rather than frost, which may be a less challenging transition to overcome physiologically (Figs. S11, S12). Furthermore, our results suggest that Rhamnaceae lineages generally retained ancestral temperate niche preferences and were more often subject to dispersal into montane tropical biomes (i.e., climatically temperate biomes in the geographical tropics) (Owens *et al*., 2017). Overall, our results do not point to dispersal dynamics as a major explanation of temperate diversity, but they do suggest that movement between temperate regions is relatively common and reflects source-sink dynamics primarily between temperate regions.

Furthermore, we found that dispersal rates out of the Mexico to Central American and Southern China hotspots were higher compared with dispersal rates from any of the other hotspots (Figs. 5, S10). These two hotspot regions cover a wide geographical distribution with high environmental heterogeneity (Figs. 2, 3, S5a), i.e., the relatively higher proportion of unique climatic hypervolumes compared to other hotspots, thus possibly providing more opportunities for exchange of lineages with subjacent regions around these two hotspots. Moreover, Mexico to Central America forms a transition zone between the Nearctic and Neotropical biotas, and more than fifty percent of Mexico is arid or semi-arid, which may have favored the dispersal of tropical lineages that were pre-adapted to survive under extreme seasonal and arid climates (Pennington & Lavin, 2016; Sosa *et al*., 2018). This suggests that the availability of diverse, heterogeneous environments may also have contributed to the evolution of Rhamnaceae species richness, and these differences may also explain differences in ‘source’ and ‘sink’ dynamics between the regions (Fig. 5).

In conclusion, our study offers an integrative approach to elucidate why certain temperate biomes, such as Mediterranean-type ecosystems, harbor high species diversity. We identify rapid *in*-*situ* diversification rates in response to the onset and expansion of seasonal biomes in the Oligocene as the best explanation; this history left a consistent signature on Rhamnaceae species composition across spatial scales, both within the temperate biomes as a whole and in regional species-rich hotspots. Although we identified a consistent global pattern, we also detected region-specific histories, with some areas illustrating higher historical connectivity through lineage dispersals to other temperate and tropical systems (e.g., Southern China, Mexico to Central America) and others indicative of evolution *in situ* as reflects their spatial isolation from other hotspots (e.g., South African Cape). Finally, our delineated species-rich regions broadly overlap with established biodiversity hotspots (Myers *et al*., 2000; CEPF, 2016) except for Southern China, which was mostly unique to Rhamnaceae. However, Southern China features exceptional plant endemism across diverse lineages, which appears to have arisen from differing mechanisms and is under increasing human threat (López-Pujol *et al*., 2011; Wang *et al*., 2022).

## Acknowledgements

This research was supported by the National Natural Science Foundation of China, key international (regional) cooperative research project (No. 31720103903), the Strategic Priority Research Program of Chinese Academy of Sciences (No. XDB31000000), the Science and Technology Basic Resources Investigation Program of China (No. 2019FY100900), the National Natural Science Foundation of China (No. 31270274), the Yunling International High-end Experts Program of Yunnan Province, China (No. YNQR-GDWG-2017-002 and No. YNQR-GDWG-2018-012), the China Scholarship Council (202004910775), the CAS President’s International Fellowship Initiative (No. 2020PB0009), the China Postdoctoral Science Foundation (CPSF), the International Postdoctoral Exchange Program, the CAS Special Research Assistant Project, a German Research Foundation grant to iDiv (DFG FZT 118, 202548816, acknowledged by REO), and the US Department of Energy (grant number DE-SC0018247 to PS, RG and DS). We are grateful to the following institutions for providing specimens or silica-dried materials: Harvard University Herbaria (A); State Herbarium of South Australia (AD); Queensland Herbarium (BRI); the herbarium of the California Academy of Sciences (CAS); the Field Museum Herbarium (F); the herbarium of Kunming Institute of Botany, Chinese Academy of Sciences (KUN); the Germplasm Bank of Wild Species and Molecular Biology Experiment Center, Kunming Institute of Botany, Chinese Academy of Sciences; National Herbarium of Victoria (MEL); the Missouri Botanical Garden Herbarium (MO); the New York Botanical Garden Herbarium (NY); the Ohio State University Herbarium (OS); Western Australian Herbarium (PERTH); the University of Texas Herbarium (TEX); and the United States National Herbarium (US). We are also grateful to Jiajin Wu for help with sampling and DNA extraction; to Ruijiang Wang for silica-dried materials of *Fenghwaia gardeniicarpa*; to Wubin Xu, Yahuang Luo, Tobias Nicolas Rojas, Friederike J. R. Wölke, Orlando Schwery, Lin Bai, Guangfu Zhu for their discussion and necessary assistance; and to the iFlora High Performance Computing Center of Germplasm Bank of Wild Species (iFlora HPC Center of GBOWS, KIB, CAS) for computing.

## Competing interests

The authors declare no conflict of interest.

## Author contributions

T.S.Y., R.E.O. and Q.T. designed the study; T.S.Y. and Q.T. collected and prepared samples for sequencing with contributions from all authors; Q.T. performed data analysis; Q.T., R.E.O. and T.S.Y. prepared the first manuscript; all authors substantially contributed to the final manuscript.

## Data availability

All sequences, occurrence and climate data used in this study will be available in the public sources and our supporting information.

## Supporting Information

**Method S1** Description and placement of fossil calibrations

**Dataset S1** The sampled species and collection information.

**Dataset S2** The concatenated alignment supermatrix of 89 low-copy loci.

**Dataset S3** The occurrence data of 1022 Rhamnaceae species from GBIF.

**Dataset S4** Information of species richness, 35 climatic variables, and classification as temperate vs. tropical biomes within each grid cell of 1° × 1°.

**Dataset S5** Classification information of the temperate vs. tropical of 1022 Rhamnaceae species.

**Fig. S1** Maximum likelihood (ML) tree of Rhamnaceae and outgroups inferred by RAxML based on the concatenated supermatrix including 89 low-copy loci under an unpartitioned GTR-GAMMA model.

**Fig. S2** Species tree of Rhamnaceae and outgroups inferred by ASTRAL-III based on 89 low-copy gene trees.

**Fig. S3** Divergence times of Rhamnaceae estimated from the concatenated supermatrix of 89 low-copy loci using treePL.

**Fig. S4** Seven diversity hotspots of Rhamnaceae.

**Fig. S5** The occupancy of hypervolume environmental space in Rhamnaceae temperate and tropical biomes.

**Fig. S6** Ancestral range reconstruction and per-area probabilities of Rhamnaceae hotspots under the BAYAREALIKE + J model using BioGeoBEARS.

**Fig. S7** Ancestral range reconstruction and per-area probabilities of Rhamnaceae buffer-hotspots under the BAYAREALIKE model using BioGeoBEARS.

**Fig. S8** Lineage through time across each Rhamnaceae hotspot using biogeographical stochastic mappings (BSMs) analysis.

**Fig. S9** Diversification rates across biomes and buffer-hotspots in Rhamnaceae.

**Fig. S10** Source and sink dynamics of Rhamnaceae in temperate regions.

**Fig. S11** Ancestral state estimates and per-state probabilities of Rhamnaceae temperate and tropical biomes (according to the geographic definition) from the best-fitting GeoSSE model. **Fig. S12** Ancestral state estimates and per-state probabilities of Rhamnaceae temperate and tropical biomes (according to the climatic definition) from the best-fitting GeoSSE model.

**Fig. S13** Global pattern of net relatedness index (NRI) plotted in grid cells of 1° × 1°.

**Table S1** The overlap percentage of climatic hypervolumes among seven hotspots.

**Table S2** Models test in BioGeoBEARS.

**Table S3** Models test in GeoSSE.

## Notes

### Competing Interest Statement

The authors have declared no competing interest.

