## Supporting Information for "Rapid *in*-*situ* diversification rates in Rhamnaceae explain the parallel evolution of high diversity in temperate biomes from global to local scales"

The following Supporting Information is available for this article:

**Method S1** Description and placement of fossil calibrations

**Dataset S1** The sampled species and collection information.

**Dataset S2** The concatenated alignment supermatrix of 89 low-copy loci.

**Dataset S3** The occurrence data of 1022 Rhamnaceae species from GBIF.

**Dataset S4** Information of species richness, 35 climatic variables, and classification as temperate vs. tropical biomes within each grid cell of 1° × 1°.

**Dataset S5** Classification information of the temperate vs. tropical of 1022 Rhamnaceae species.

**Fig. S1** Maximum likelihood (ML) tree of Rhamnaceae and outgroups inferred by RAxML based on the concatenated supermatrix including 89 low-copy loci under an unpartitioned GTR-GAMMA model.

**Fig. S2** Species tree of Rhamnaceae and outgroups inferred by ASTRAL-III based on 89 low-copy gene trees.

**Fig. S3** Divergence times of Rhamnaceae estimated from the concatenated supermatrix of 89 low-copy loci using treePL.

**Fig. S4** Seven diversity hotspots of Rhamnaceae.

**Fig. S5** The occupancy of hypervolume environmental space in Rhamnaceae temperate and tropical biomes.

**Fig. S6** Ancestral range reconstruction and per-area probabilities of Rhamnaceae hotspots under the BAYAREALIKE + J model using BioGeoBEARS.

**Fig. S7** Ancestral range reconstruction and per-area probabilities of Rhamnaceae buffer-hotspots under the BAYAREALIKE model using BioGeoBEARS.

**Fig. S8** Lineage through time across each Rhamnaceae hotspot using biogeographical stochastic mappings (BSMs) analysis.

**Fig. S9** Diversification rates across biomes and buffer-hotspots in Rhamnaceae.

**Fig. S10** Source and sink dynamics of Rhamnaceae in temperate regions.

**Fig. S11** Ancestral state estimates and per-state probabilities of Rhamnaceae temperate and tropical biomes (according to the geographic definition) from the best-fitting GeoSSE model.

**Fig. S12** Ancestral state estimates and per-state probabilities of Rhamnaceae temperate and tropical biomes (according to the climatic definition) from the best-fitting GeoSSE model.

**Fig. S13** Global pattern of net relatedness index (NRI) plotted in grid cells of 1° × 1°.

**Table S1** The overlap percentage of climatic hypervolumes among seven hotspots.

**Table S2** Models test in BioGeoBEARS.

**Table S3** Models test in GeoSSE.

**Method S1** Description and placement of fossil calibrations. SG, stem group; CG, crown group.

Rhamnaceae fossils

A. **Node: SG Rhamnaceae**

Secondary calibration.

**Reference:** Onstein *et al.* (2016); Hauenschild *et al.* (2018).

**Calibration:** Most recently estimated Rhamnaceae stem ages were roughly equal to or greater than 100 Ma, such as c. 98.7 Ma (88.6–113.7 Ma; 95% HPD) (Onstein *et al.*, 2016), ca. 100.0–104.3 Ma (Hauenschild *et al.*, 2018). Integrating previous age estimates, we applied 100.0 Ma and 113.7 Ma as the minimum-and maximum-age calibrations of the stem age of Rhamnaceae.

B. **Node: CG Rhamnaceae**

**Fossils:** Flowers of *Coahuilanthus belindae* from the late Cretaceous (late Campanian) of Cerro del Pueblo Formation, Coahuila, Mexico (Calvillo-Canadell & Cevallos-Ferriz, 2007).

**Reference:** Calvillo-Canadell & Cevallos-Ferriz (2007).

**Calibration:** The *Coahuilanthus belindae* is the earliest reliable fossil of Rhamnaceae. The floral morphology of *Coahuilanthus belindae* is closely related to extant Rhamneae (e.g., *Rhamnu*s, *Sageretia* and *Berchemia*), based on floral cup structures and parts of the perianth (e.g. five acute, triangular, slightly keeled sepals). We therefore placed it at the crown of the Rhamnaceae and considered two boundaries of late Campanian (72.1–83.6 Ma) as the minimum- and maximum-age calibrations.

C. **Node: CG Rhamneae**

**Fossils:** The leaves of *Berhamniphyllum* from the late Cretaceous (Maastrichtian) of Guaduas Formation, central Colombia (Correa *et al.*, 2010).

**Reference:** Correa et al. (2010).

**Calibration:** The leaves of *Berhamniphyllum* from Colombia (ca. 68 Ma) resemble an extinction clade of the tribe Rhamneae (e.g., *Berchemia* and *Rhamnidium* with in tribe Rhamneae) , based on uniformly curved, eucamptodromous secondary venation and closely spaced, percurrent tertiaries oriented perpendicular to the midvein (Correa *et al.*, 2010; Zhou *et al.*, 2019). We therefore placed it at the crown of Rhamneae (68 Ma).

D. **Node: CG Paliureae**

**Fossils:** The winged fruit of *Archaeopaliurus boyacensis* from the late Cretaceous (Maastrichtian) of Guaduas Formation, central Colombia (Correa *et al.*, 2010).

**Reference:** Correa et al. (2010)

**Calibration:** The *Archaeopaliurus boyacensis* has been reported from Colombia (ca. 68 Ma) (Correa et al. 2010). This fossil has winged fruit, which is closely related to extant *Paliurus* fruits. We therefore placed it at the crown of Paliureae.

E. **Node: SG *Berchemia***

**Fossils:** An endocarp of *Berchemia mellerae* from the Middle Eocene of the Messel Fromation, Germany (Collinson et al. 2012).

**Reference:** Collinson et al. 2012.

**Calibration:** *Berchemia mellerae* is the oldest fossil (endocarp) ever found in *Berchemia* and is dated to the middle Eocene (ca. 47 Ma) (Collinson *et al.*, 2012; Zhou *et al.*, 2019). We therefore placed it at the stem of *Berchemia*.

F. **Node: CG Ventilageneae**

**Fossils:** Fruits of *Ventilago tibetensis* from the middle Eocene (ca. 47 Ma) of Niubao Formation, Jianglang, central Tibetan Plateau (Del Rio *et al.*, 2021).

**Reference:** Del Rio et al. (2021).

**Calibration:** *Ventilago tibetensis* shows similarities with the sister genera *Smythea* and *Ventilago*, based on several well-preserved fossils fruits with an apically extended elongate wing and a conical hypanthium enclosing two-thirds of an elliptical and apically pointed seed chamber. *Ventilago tibetensis* is the earliest fossil record of the tribe Ventilagineae (Del Rio et al., 2021). Therefore, we place it at the crown of Ventilageneae (47 Ma).

G. **Node: CG *Paliurus***

**Fossils:** Fruits of the *Paliurus clarnensis* from the middle Eocene of the Clarno Formation, Oregon (Burge & Manchester, 2008).

**Reference:** Burge and Manchester (2008).

**Calibration:** *Paliurus clarnensis* is the reliable member of *Paliurus* and is dated to the middle Eocene (ca. 44 Ma). The characters of these fossil fruits largely match the extant members of the genus. We therefore placed it at the crown of *Paliurus* (44 Ma).

**Fig. S1** Maximum likelihood (ML) tree of Rhamnaceae and outgroups inferred by RAxML based on the concatenated supermatrix of 89 low-copy loci under an unpartitioned GTR-GAMMA model. Branch lengths are proportional to per-site substitution rate. Branch labels are bootstrap support values. The tree has been broken into panels for display purposes; bold letters indicate where branches attach across panels.

**Fig. S2** Species tree of Rhamnaceae and outgroups inferred in ASTRAL-III based on 89 gene trees. Branch lengths are coalescent units measured in ASTRAL-III for internal branches and not terminal branches (branch lengths of terminal branches are therefore arbitrary and meaningless). Branch labels are local posterior probability values. The tree has been broken into panels for display purposes; bold letters indicate where branches attach across panels.

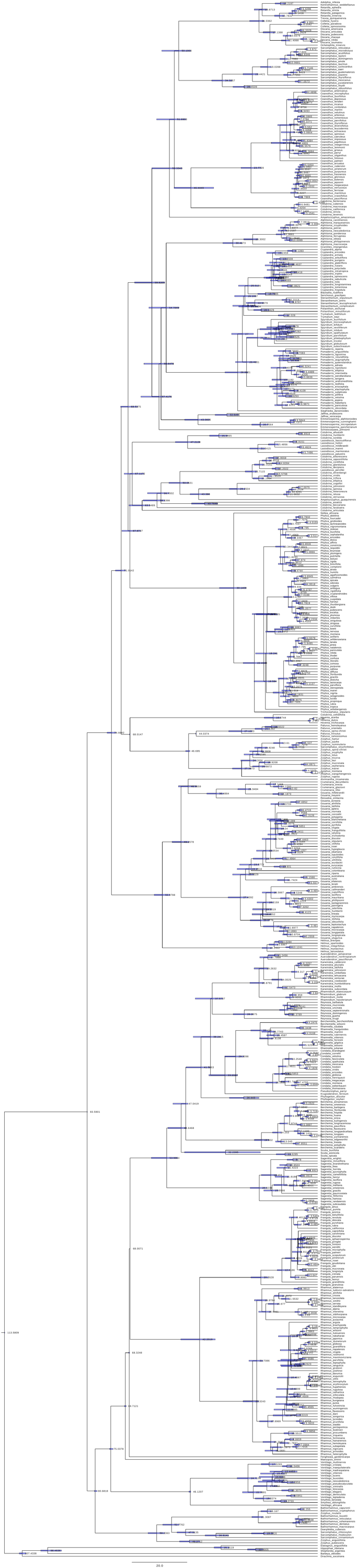

**Fig. S3** Divergence times of Rhamnaceae estimated from the concatenated supermatrix of 89 low-copy loci using treePL. The numbers represent the estimated median age of nodes. The blue bars show the range of the upper and lower confidence intervals calculated from 100 bootstrap replications.

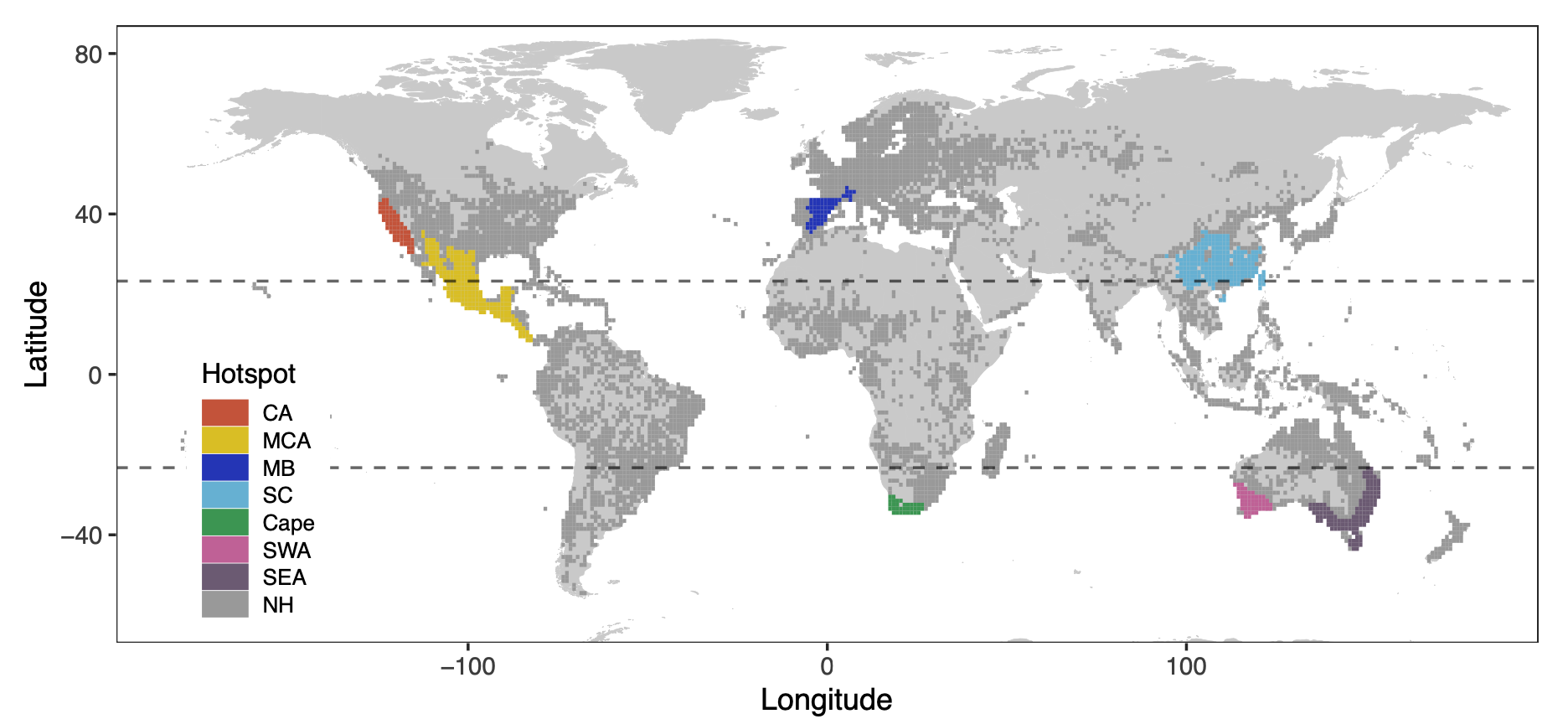

**Fig. S4** Seven Rhamnaceae diversity hotspots inferred by the Hotspot Analysis Plugin in QGIS3, colored grid cells with Z scores ≥ 1.65 are hotspots. CA, California; MCA, Mexico to Central America; MB, Mediterranean Basin; SC, Southern China; SWE, Southwest Australia; SEA, Southeast Australia; NH, non-hotspot.

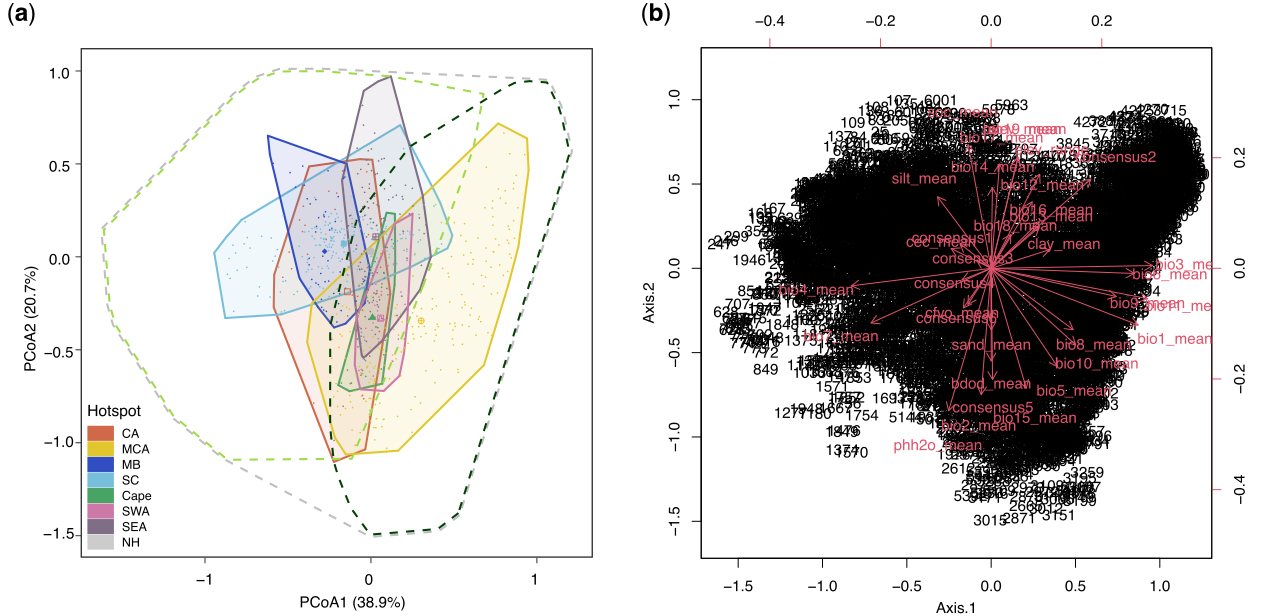

**Fig. S5** The occupancy and biplot of hypervolume environmental space in Rhamnaceae temperate and tropical biomes. (a) Colored polygons are two-dimensional environmental hypervolumes based on 772 assemblages (grid cells) in seven hotspots using principal coordinates analysis (PCoA). The light green, dark green and gray dashed polygons are the environmental hypervolumes of respectively temperate, tropical (according to the climatic definition) and non-hotspot regions. Colored points represent grid cells. Colored shapes are hypervolume centroids. CA, California; MCA, Mexico to Central America; MB, Mediterranean Basin; SC, Southern China; SWE, Southwest Australia; SEA, Southeast Australia. (b) The biplot of hypervolume climate space in Rhamnaceae. The number is the name of each grid cell.

**Fig. S6** Ancestral range estimates and per-area probabilities of Rhamnaceae hotspots under the BAYAREALIKE + J model using BioGeoBEARS. Boxes at each node are color-coded for the area with the probability of each area. CA, California; MCA, Mexico to Central America; MB, Mediterranean Basin; SC, Southern China; SWE, Southwest Australia; SEA, Southeast Australia; NH, non-hotspot.

**Fig. S7** Ancestral range eatimates and per-area probabilities of Rhamnaceae buffer-hotspots under the BAYAREALIKE model using BioGeoBEARS. Boxes at each node are color-coded for the area with the probability of each area. CA, California; MCA, Mexico to Central America; MB, Mediterranean Basin; SC, Southern China; SWE, Southwest Australia; SEA, Southeast Australia; NH, non-hotspot.

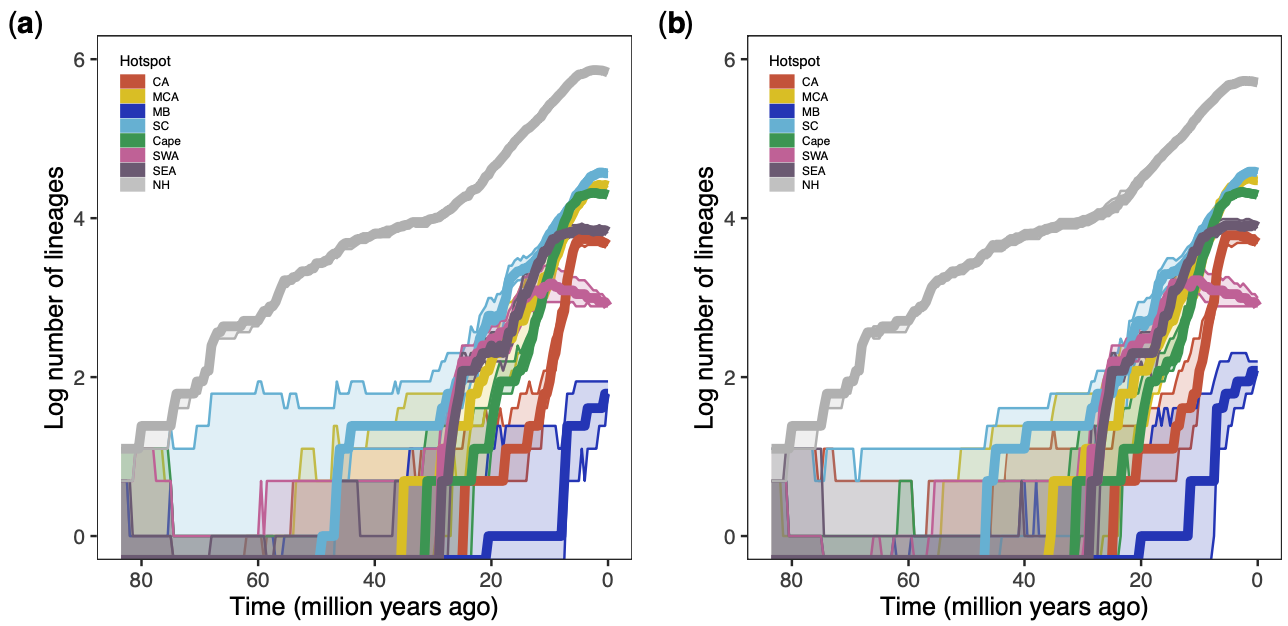

**Fig. S8** Lineage through time (LTT) across each Rhamnaceae hotspot. LTT plot across (a) each hotspot and non-hotspot, (b) buffer-hotspot and non-hotspot from 100 biogeographical stochastic mappings (BSMs) analysis using a sliding window analysis and median value (bold color lines) from 95% confidence interval (solid color areas) of the number of lineages on a logarithmic scale occupying each region at each time bin. CA, California; MCA, Mexico to Central America; MB, Mediterranean Basin; SC, Southern China; SWE, Southwest Australia; SEA, Southeast Australia; NH, non-hotspot.

**Fig. S9** Diversification rates across biomes in Rhamnaceae. Estimated parameter distributions in temperate vs. tropical biomes (according to the climatic definition) as well as of Rhamnaceae lineages evolving in five buffer-hotspots. The Mediterranean Basin and the Southwest Australian were excluded because sampling of species in these two hotspots was less than 10% compared to elsewhere. Boxplots showing (a) speciation rates (lineage/Myr), (b) extinction rates (lineage/Myr) and (c) net diversification rates (lineage/Myr); (d) speciation rates (lineage/Myr), (e) extinction rates (lineage/Myr) and (f) net diversification rates (lineage/Myr) of the five buffer-hotspots. Boxes are colored according to the region, and represent parameter distributions from the Bayesian MCMC using the best-fitting GeoSSE model on the time-calibrated Rhamnaceae phylogenetic tree. Dashed lines show in figure (a-c) averaged parameter of non-hotspot lineages. CA, buffer-California; MCA, buffer-Mexico to Central America; SC, buffer-Southern China; Cape, buffer-Cape; SEA, buffer-Southeast Australia.

**Fig. S10** Source and sink dynamics of Rhamnaceae in temperate regions. Boxes are colored according to temperate biomes (according to the climatic definition) and buffer-hotspots, and represent parameter distributions (dispersal rates out of a region minus dispersal rates into that region) from the Bayesian MCMC from the best-fitting GeoSSE model on the time-calibrated Rhamnaceae phylogenetic tree. CA, buffer-California; MCA, buffer-Mexico to Central America; SC, buffer-Southern China; Cape, buffer-Cape; SEA, buffer-Southeast Australia.

**Fig. S11** Ancestral state estimates and per-state probabilities of Rhamnaceae biomes (according to the geographic definition) from the best-fitting GeoSSE model.

**Fig. S12** Ancestral state estimates and per-state probabilities of Rhamnaceae biomes (according to the climatic definition) from the best-fitting GeoSSE model.

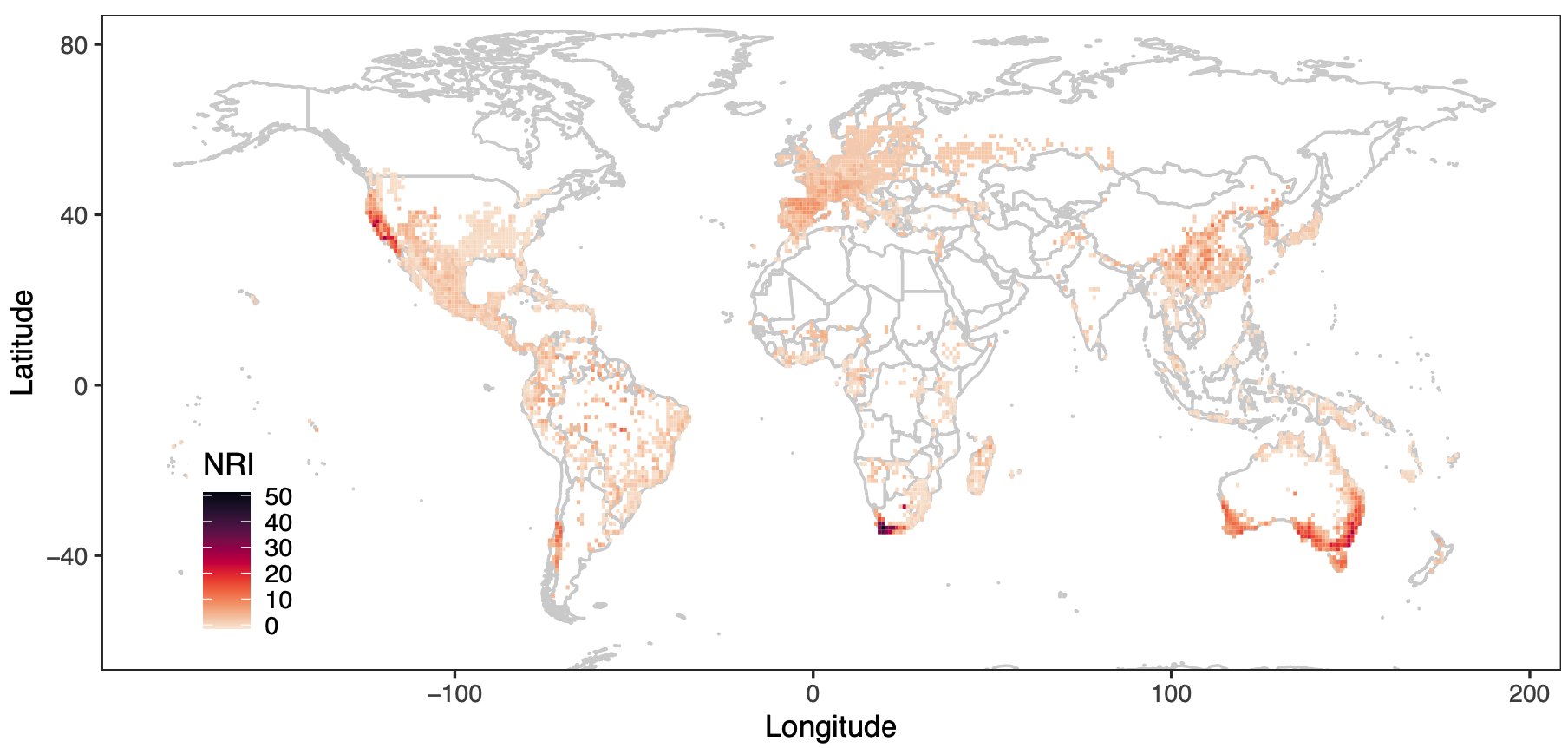

**Fig. S13** Global Net Relatedness Index (NRI) map plotted in grid cells of 1° × 1°.

**Table S1** The overlap percentage of climatic hypervolumes among seven hotspots

|  | CA | MCA | MB | SC | Cape | SWA | SEA | Average |
| --- | --- | --- | --- | --- | --- | --- | --- | --- |
| CA | NA | 45.6% | 37.7% | 50.0% | 15.8% | 12.1% | 24.9% | 31.0% |
| MCA | 26.9% | NA | 5.2% | 12.7% | 11.6% | 16.8% | 15.8% | 14.8% |
| MB | 79.8% | 18.7% | NA | 72.6% | 0.5% | 0.2% | 15.2% | 31.2% |
| SC | 46.7% | 20.1% | 32.0% | NA | 4.5% | 5.6% | 38.9% | 24.7% |
| Cape | 77.3% | 96.4% | 1.1% | 23.8% | NA | 75.5% | 64.1% | 56.4% |
| SWA | 45.1% | 100.0% | 0.3% | 22.4% | 57.7% | NA | 64.0% | 48.2% |
| SEA | 38.3% | 43.7% | 11.0% | 30.3% | 20.1% | 26.3% | NA | 28.3% |

Note: CA, California; MCA, Mexico to Central America; MB, Mediterranean Basin; SC, Southern China; SWE, Southwest Australia; SEA, Southeast Australia.

Read from left to right, e.g., the California is for 45.6% overlap compared to Mexico to Certral America, whereas Mexico to Central America is for 26.9% overlap compared to California. The average of overlap percentage of California with other six hotspots is 31%.

**Table S2** Models test in BioGeoBEARS.

| Region | Model | LnL | numparams | d | e | j | AICc | AICc_wt |
| --- | --- | --- | --- | --- | --- | --- | --- | --- |
| hotspots | DEC | -934.2 | 2 | 0.0031 | 0.025 | 0 | 1872 | 2.7e-58 |
|  | DEC+J | -931.4 | 3 | 0.0027 | 0.015 | 0.0026 | 1869 | 1.6e-57 |
|  | DIVALIKE | -986.5 | 2 | 0.0045 | 0.066 | 0 | 1977 | 4.9e-81 |
|  | DIVALIKE+J | -976.7 | 3 | 0.0034 | 0.037 | 0.0045 | 1959 | 3.5e-77 |
|  | BAYAREALIKE | -803.6 | 2 | 0.0017 | 0.053 | 0 | 1611 | 0.14 |
|  | **BAYAREALIKE+J** | **-800.8** | **3** | **0.0016** | **0.048** | **0.0014** | **1608** | **0.86** |
| buffer-hotspots | DEC | -846.5 | 2 | 0.0029 | 0.030 | 0 | 1697 | 1.9e-31 |
|  | DEC+J | -842.8 | 3 | 0.0022 | 0.015 | 0.0034 | 1692 | 2.9e-30 |
|  | DIVALIKE | -889.3 | 2 | 0.0041 | 0.075 | 0 | 1783 | 4.7e-50 |
|  | DIVALIKE+J | -878.4 | 3 | 0.0028 | 0.037 | 0.0051 | 1763 | 9.1e-46 |
|  | **BAYAREALIKE** | **-776.1** | **2** | **0.0022** | **0.067** | **0** | **1556** | **0.71** |
|  | BAYAREALIKE+J | -776 | 3 | 0.0021 | 0.066 | 0.0003 | 1558 | 0.29 |

Note: ‘d’ refers to ‘dispersal’, ‘e’ to ‘extinction rate’and ‘j’ to ‘jump dispersal’.

LnL=log marginal likelihood, AICc=corrected Akaike, AICc_wt=weighted AICc.

**Table S3** Models test in GeoSSE.

| Region | Model | Df | lnLik | AIC | ChiSq | Pr(>\|Chi\|) |
| --- | --- | --- | --- | --- | --- | --- |
| California | full | 7 | -2190.4 | 4394.8 |  |  |
|  | equal.d | 6 | -2204.1 | 4420.2 | 27.347 | 1,70E-04 *** |
|  | equal.s | 5 | -2194.4 | 4398.9 | 8.060 | 0.01777 * |
|  | **equal.x** | **6** | **-2190.6** | **4393.3** | **0.436** | **0.50895** |
|  | equal.sxd | 3 | -2207.4 | 4420.8 | 34.012 | 7,41E-04 *** |
|  | equal.sx | 4 | -2194.6 | 4397.3 | 8.447 | 0.03762 * |
|  | equal.sd | 4 | -2207.4 | 4422.7 | 33.917 | 2,06E-04 *** |
|  | equal.xd | 5 | -2204.1 | 4418.2 | 27.346 | 1,15E-03 *** |
| Mexico to Central America | full | 7 | -2337.8 | 4689.7 |  |  |
|  | equal.d | 6 | -2357.6 | 4727.2 | 39.500 | 3,28E-07 *** |
|  | equal.s | 5 | -2342.4 | 4694.8 | 9.085 | 0.010647 * |
|  | **equal.x** | **6** | **-2338.2** | **4688.3** | **0.617** | **0.432120** |
|  | equal.sxd | 3 | -2362.7 | 4731.3 | 49.622 | 4,33E-07 *** |
|  | equal.sx | 4 | -2343.7 | 4695.5 | 11.759 | 0.008258 ** |
|  | equal.sd | 4 | -2358.9 | 4725.8 | 42.124 | 3,78E-06 *** |
|  | equal.xd | 5 | -2359.1 | 4728.2 | 42.457 | 6,03E-07 *** |
| Southern China | full | 7 | -2325.4 | 4664.8 |  |  |
|  | equal.d | 6 | -2373.9 | 4759.8 | 97.048 | <2e-16 *** |
|  | **equal.s** | **5** | **-2325.6** | **4661.1** | **0.331** | **0.8474** |
|  | equal.x | 6 | -2325.8 | 4663.7 | 0.891 | 0.3451 |
|  | equal.sxd | 3 | -2382.3 | 4770.6 | 113.859 | <2e-16 *** |
|  | equal.sx | 4 | -2328.1 | 4664.2 | 5.388 | 0.1455 |
|  | equal.sd | 4 | -2374.5 | 4757.1 | 98.305 | <2e-16 *** |
|  | equal.xd | 5 | -2381.1 | 4772.2 | 111.453 | <2e-16 *** |
| Cape | full | 7 | -2178.0 | 4370.0 |  |  |
|  | equal.d | 6 | -2199.5 | 4411.0 | 42.944 | 5,63E-08 *** |
|  | equal.s | 5 | -2184.2 | 4378.3 | 12.258 | 0.0021792 ** |
|  | **equal.x** | **6** | **-2178.1** | **4368.1** | **0.086** | **0.7687935** |
|  | equal.sxd | 3 | -2208.4 | 4422.8 | 60.735 | 2,03E-09 *** |
|  | equal.sx | 4 | -2187.1 | 4382.3 | 18.231 | 0.0003941 *** |
|  | equal.sd | 4 | -2208.1 | 4424.1 | 60.070 | 5,68E-10 *** |
|  | equal.xd | 5 | -2199.5 | 4409.0 | 42.941 | 4,74E-07 *** |
| Southeast Australia | full | 7 | -2214.6 | 4443.2 |  |  |
|  | equal.d | 6 | -2232.1 | 4476.2 | 35.084 | 3,16E-06 *** |
|  | equal.s | 5 | -2219.8 | 4449.7 | 10.537 | 0.005152 ** |
|  | **equal.x** | **6** | **-2214.6** | **4441.2** | **0.050** | **0.823240** |
|  | equal.sxd | 3 | -2236.4 | 4478.7 | 43.565 | 7,90E-06 *** |
|  | equal.sx | 4 | -2220.5 | 4449.0 | 11.818 | 0.008034 ** |
|  | equal.sd | 4 | -2235.9 | 4479.8 | 42.608 | 2,98E-06 *** |
|  | equal.xd | 5 | -2232.1 | 4474.2 | 35.084 | 2,41E-05 *** |
| Combined seven hotspots | **full** | **7** | **-2585.0** | **5184.0** |  |  |
|  | equal.d | 6 | -2599.7 | 5211.5 | 29.520 | 5,53E-05 *** |
|  | equal.s | 5 | -2594.7 | 5199.5 | 19.523 | 5,76E-02 *** |
|  | equal.x | 6 | -2587.1 | 5186.1 | 4.146 | 0.04174 * |
|  | equal.sxd | 3 | -2624.1 | 5254.2 | 78.281 | 4,44E-13 *** |
|  | equal.sx | 4 | -2617.7 | 5243.5 | 65.505 | 3,91E-11 *** |
|  | equal.sd | 4 | -2617.2 | 5242.4 | 64.472 | 6,51E-11 *** |
|  | equal.xd | 5 | -2599.7 | 5209.5 | 29.519 | 3,89E-04 *** |
| buffer-California | full | 7 | -2193.8 | 4401.5 |  |  |
|  | equal.d | 6 | -2203.7 | 4419.4 | 199.109 | 8,11E-03 *** |
|  | equal.s | 5 | -2197.9 | 4405.7 | 82.158 | 0.01644 * |
|  | **equal.x** | **6** | **-2193.9** | **4399.8** | **0.2984** | **0.58486** |
|  | equal.sxd | 3 | -2207.2 | 4420.4 | 269.127 | 2,07E-02 *** |
|  | equal.sx | 4 | -2197.9 | 4403.7 | 82.169 | 0.04174 * |
|  | equal.sd | 4 | -2207.2 | 4422.4 | 269.120 | 6,14E-03 *** |
|  | equal.xd | 5 | -2203.7 | 4417.4 | 199.109 | 4,75E-02 *** |
| buffer-Mexico to Central America | full | 7 | -2336.8 | 4687.5 |  |  |
|  | equal.d | 6 | -2347.2 | 4706.4 | 209.008 | 4,84E-03 *** |
|  | **equal.s** | **5** | **-2337.6** | **4685.1** | **15.849** | **0.45273** |
|  | **equal.x** | **6** | **-2336.9** | **4685.9** | **0.3130** | **0.57586** |
|  | equal.sxd | 3 | -2349.8 | 4705.5 | 259.728 | 3,21E-02 *** |
|  | equal.sx | 4 | -2341.2 | 4690.4 | 88.622 | 0.03118 * |
|  | equal.sd | 4 | -2348.6 | 4705.1 | 235.871 | 3,05E-02 *** |
|  | equal.xd | 5 | -2347.5 | 4705.0 | 214.667 | 2,18E-02 *** |
| buffer-Southern China | full | 7 | -2309.6 | 4633.3 |  |  |
|  | equal.d | 6 | -2343.0 | 4698.1 | 66.812 | 3,33E-13 *** |
|  | **equal.s** | **5** | **-2310.2** | **4630.3** | **1.051** | **0.591119** |
|  | equal.x | 6 | -2312.3 | 4636.5 | 5.258 | 0.021852 * |
|  | equal.sxd | 3 | -2349.5 | 4705.1 | 79.780 | 2,22E-13 *** |
|  | equal.sx | 4 | -2315.6 | 4639.1 | 11.833 | 0.007978 ** |
|  | equal.sd | 4 | -2345.0 | 4697.9 | 70.670 | 3,11E-12 *** |
|  | equal.xd | 5 | -2348.1 | 4706.1 | 76.873 | < 2.2e-16 *** |
| buffer-Cape | full | 7 | -2172.2 | 4358.4 |  |  |
|  | equal.d | 6 | -2186.9 | 4385.8 | 29.432 | 5,79E-05 "*** " |
|  | equal.s | 5 | -2178.7 | 4367.4 | 13.057 | 0.0014612 "** " |
|  | **equal.x** | **6** | **-2172.2** | **4356.4** | **0.028** | **0.8664778** |
|  | equal.sxd | 3 | -2197.4 | 4400.8 | 50.413 | 2,96E-07 "*** " |
|  | equal.sx | 4 | -2181.8 | 4371.5 | 19.112 | 0.0002592 "*** " |
|  | equal.sd | 4 | -2196.3 | 4400.7 | 48.322 | 1,82E-07 "*** " |
|  | equal.xd | 5 | -2186.9 | 4383.8 | 29.430 | 4,07E-04 "*** " |
| buffer-Southeast Australia | full | 7 | -2218.2 | 4450.3 |  |  |
|  | equal.d | 6 | -2229.9 | 4471.7 | 23.444 | 1,29E-03 *** |
|  | equal.s | 5 | -2224.7 | 4459.5 | 13.179 | 0.001375 ** |
|  | **equal.x** | **6** | **-2218.2** | **4448.3** | **0.045** | **0.832318** |
|  | equal.sxd | 3 | -2235.3 | 4476.7 | 34.363 | 6,28E-04 *** |
|  | equal.sx | 4 | -2225.1 | 4458.2 | 13.946 | 0.002980 ** |
|  | equal.sd | 4 | -2234.7 | 4477.4 | 33.069 | 3,12E-04 *** |
|  | equal.xd | 5 | -2229.9 | 4469.7 | 23.444 | 8,11E-03 *** |
| buffer-Combined seven hotspots | **full** | **7** | **-2547.2** | **5108.4** |  |  |
|  | equal.d | 6 | -2550.5 | 5113.0 | 6.561 | 0.01042 * |
|  | equal.s | 5 | -2557.5 | 5125.0 | 20.635 | 3,31E-02 *** |
|  | equal.x | 6 | -2548.7 | 5109.3 | 2.884 | 0.08944 . |
|  | equal.sxd | 3 | -2578.6 | 5163.1 | 62.682 | 7,92E-10 *** |
|  | equal.sx | 4 | -2577.7 | 5163.4 | 60.991 | 3,61E-10 *** |
|  | equal.sd | 4 | -2565.9 | 5139.8 | 37.369 | 3,85E-05 *** |
|  | equal.xd | 5 | -2550.8 | 5111.5 | 7.109 | 0.02860 * |
| Temperate biomes (geographic definition) | full | 7 | -2415.2 | 4844.4 |  |  |
|  | **equal.d** | **6** | **-2415.2** | **4842.4** | **0.028** | **0.86785** |
|  | equal.s | 5 | -2432.8 | 4875.7 | 35.237 | 2,23E-05 *** |
|  | equal.x | 6 | -2417.3 | 4846.6 | 4.216 | 0.04005 * |
|  | equal.sxd | 3 | -2459.2 | 4924.4 | 87.988 | <2.2e-16 *** |
|  | equal.sx | 4 | -2459.1 | 4926.2 | 87.759 | <2.2e-16 *** |
|  | equal.sd | 4 | -2433.7 | 4875.4 | 36.953 | 4,71E-05 *** |
|  | equal.xd | 5 | -2417.8 | 4845.5 | 5.126 | 0.07709 . |
| Temperate biomes (climatic definition) | full | 7 | -2464.9 | 4943.8 |  |  |
|  | equal.d | 6 | -2477.5 | 4967.0 | 25.216 | 5,12E-04 *** |
|  | equal.s | 5 | -2484.8 | 4979.6 | 39.831 | 2,24E-06 *** |
|  | **equal.x** | **6** | **-2465.3** | **4942.6** | **0.836** | **0.3607** |
|  | equal.sxd | 3 | -2524.5 | 5055.1 | 119.284 | <2.2e-16 *** |
|  | equal.sx | 4 | -2515.8 | 5039.7 | 101.859 | <2.2e-17 *** |
|  | equal.sd | 4 | -2510.9 | 5029.7 | 91.943 | <2.2e-18 *** |
|  | equal.xd | 5 | -2477.5 | 4965.0 | 25.217 | 3,34E-03 *** |

Note: ‘d’ refers to ‘dispersal rate’, ‘s’ to ‘speciation rate’ and ‘x’ to ‘extinction rate’.

‘Df’=degrees of freedom, lnLik=log-likelihood, AIC=Akaike Information Criterion, ChiSq=Chi square.

Signif. codes: 0 '***' 0.001 '**' 0.01 '*' 0.05 '.' 0.1 ' ' 1
